# Computing the effects of excitatory-inhibitory balance on neuronal input-output properties

**DOI:** 10.1101/2025.03.09.642210

**Authors:** Alex Reyes

## Abstract

In sensory systems, stimuli are represented through the diverse firing responses and receptive fields of neurons. These features emerge from the interaction between excitatory (*E*) and inhibitory (*I*) neuron populations within the network. Changes in sensory inputs alter this balance, leading to shifts in firing patterns and the input-output properties of individual neurons and the network. While these phenomena have been studied extensively with experiments and theory, the underlying principles for combining *E* and *I* inputs are still unclear. Here, the rules for probabilistically combining *E* and *I* inputs are derived that describe how neurons in a feedforward inhibitory circuit respond to stimuli. This simple model is broadly applicable, capturing a wide range of response features that would otherwise require multiple separate models and offers insights into the cellular and network mechanisms influencing the input-output properties of neurons, gain modulation, and the emergence of diverse temporal firing patterns.

**Author Summary:** Sensory stimuli activate a broad network of excitatory and inhibitory neurons. The response of individual neurons is often complex and influenced by the animal’s state—such as whether it is resting, moving, or attending to a specific environmental cue. To understand how stimulus features are encoded and modulated, it is essential to examine how synaptic inputs are integrated within individual neurons and across neural networks. In this manuscript, I propose a set of rules for combining excitatory and inhibitory synaptic inputs in neural circuits. This simple, general model captures key features of sensory-driven responses.

## Introduction

A common motif in the sensory systems is the feedforward inhibitory circuit, where excitatory afferents from an external source synapse onto both excitatory (*E*) and inhibitory (*I*) neurons, with inhibitory neurons then synapsing back onto *E* cells [1, 2, 3, 4, 5]. During stimulation, interactions within this circuit generate complex dynamics and shape the receptive field properties of neurons. As stimulus parameters vary, the balance between *E* and *I* inputs shifts, leading to both qualitative and quantitative changes in neuronal sensitivity and evoked firing patterns [1, 4, 6, 7].

A neuron’s response to a stimulus is characterized by its input–output (I–O) curve, where the input typically refers to synaptic current and the output to firing rate. Compared to neurons *in vitro* [8, 9], neurons *in vivo* exhibit more diverse I-O profiles. Background excitation and inhibition from ongoing network activity introduce membrane potential fluctuations, allowing neurons to respond even to weak inputs that would otherwise remain subthreshold, and to generate smoothly increasing I–O curves [10, 11, 12, 13]. In some cases, responses initially increase with stimulus intensity but then decline after reaching a peak [14, 15, 16, 17], likely due to strong inhibitory recruitment [18, 19]. I–O curves also depend stimulus duration: brief stimuli (5–50 ms) evoke responses sensitive to the timing of excitation and inhibition [2, 4, 20, 21], while longer stimuli (hundreds of milliseconds) evoke responses that vary with the average synaptic current generated during barrages [22, 23, 24].

Moreover, the I-O curves may change depending on the animal’s state, such as when it is at rest, in motion [25, 26] or actively attending [27, 28]. To maintain selectivity to stimulus features across states, the slope of the I-O curve should change without affecting the minimum input needed to evoke firing [13, 29]. This multiplicative (or divisive) gain modulation can arise from feedback from neighboring *E* and/or *I* neurons [30, 31, 32, 33, 34], feedforward inhibition [21] or the combination of synaptic noise and conductance [10]. Additive (or subtractive) modulation, by contrast, shifts the activation threshold without altering gain, thereby affecting tuning curve width [13].

Finally, differences in *E* -*I* balance can produce diverse temporal firing responses. Some neurons exhibit continuous firing throughout the stimulus duration, while others fire transiently at the stimulus onset [16, 22, 35, 36, 37, 38, 39] and/or at the offset [39, 40, 41, 42] . These firing profiles are observed in cortical and subcortical neurons [42] and may be generated locally [22, 41, 42, 43, 44] or inherited from upstream sources [40]. Additionally, a neuron’s response type may change based on the stimulus intensity [16] or whether the preferred stimulus is presented [35, 38, 45, 46].

Identifying common operating principles across these phenomena will provide valuable insights into potential mechanisms. This study aims to derive rules for calculating *E-I* balance in feed-forward inhibitory circuits. The model combines *E* and *I* inputs probabilistically and links the associated changes in responses and I-O curves to the synaptic and network properties. The model reproduces and clarifies the conditions for gain modulation, non-monotonic I-O curves, and diverse firing patterns.

## Results

The following sections begin with an idealized feedforward inhibitory network to introduce core concepts of the probabilistic interaction between excitatory and inhibitory inputs. These principles are then extended to more physiologically realistic conditions that include multiple, temporally distributed inputs. Finally, the model is applied to examine neuronal input–output properties, such as gain modulation and temporal firing profiles.

### Simple model

Consider a hypothetical circuit consisting of a postsynaptic excitatory neuron (henceforth termed the “reference cell”) and a single inhibitory neuron, both receiving an excitatory postsynaptic potential (EPSP) from a common external afferent (Fig. 1A). During stimulation, the afferent evokes an EPSP in both cells with probability *p*_*E*_. The firing of the *I* cell—and consequently the occurrence of an inhibitory postsynaptic potential (IPSP) in the *E* cell—requires that an EPSP first arise in the *I* neuron. If that EPSP is suprathreshold, the *I* cell fires with conditional probability

**Figure 1.**
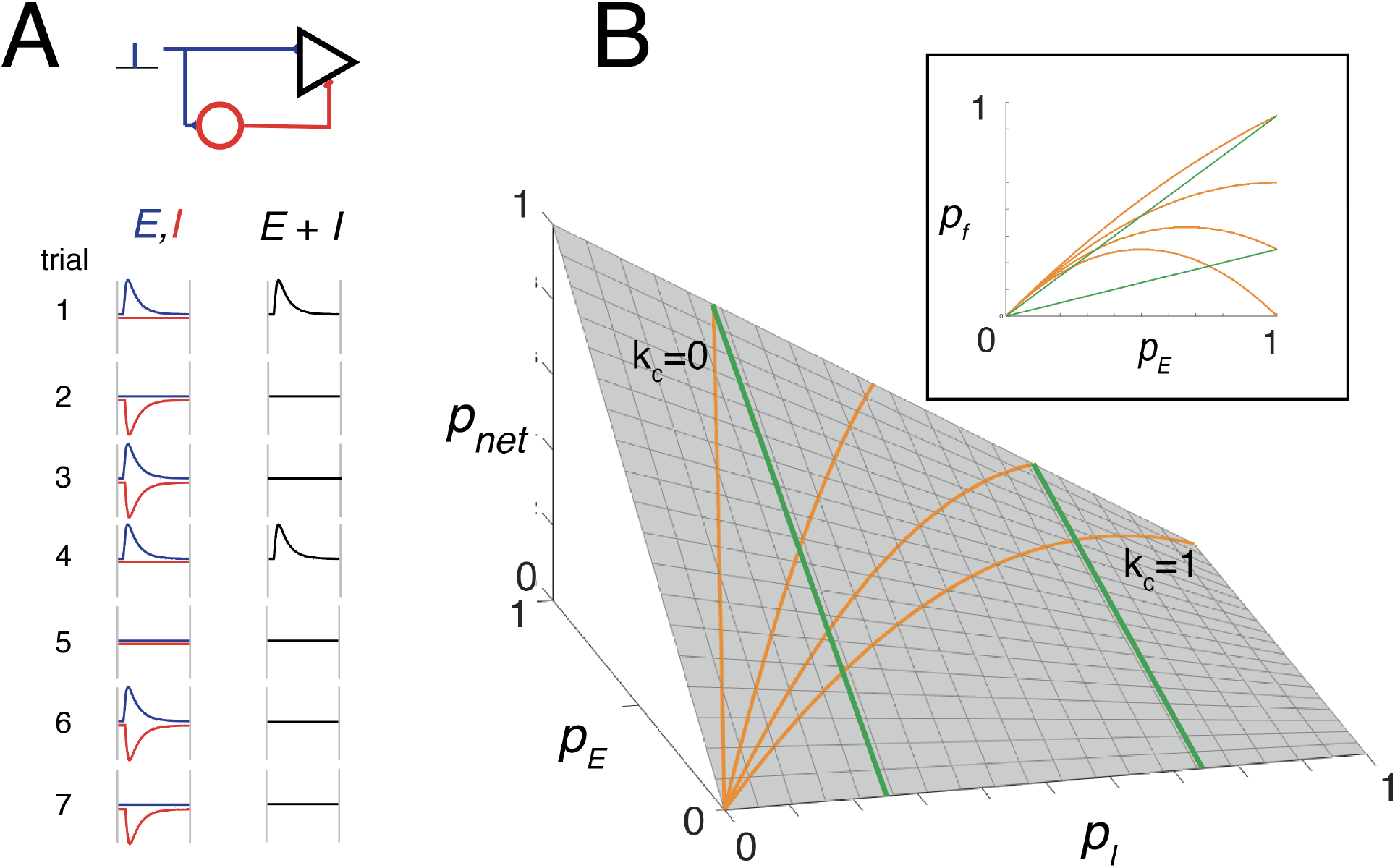
Toy model illustrating the basic principles. **A, top**, schematic of a simple feedforward inhibitory circuit. A postsynaptic excitatory neuron (black triangle) and a local inhibitory interneuron (red circle) both receive a single excitatory input from an external afferent. An IPSP is generated in the postsynaptic *E* neuron whenever the interneuron fires. **Bottom**, EPSPs (blue) and IPSPs (red) recorded in the postsynaptic *E* neuron over seven stimulus trials. An EPSP is shown as canceled when it coincides with an IPSP, but more generally Eq. 1 describes the probabilistic outcome: over *n* trials, the expected number of uncanceled EPSPs is *np*_net_. **B**, surface plot of the net probability *p*_*net*_ as a function of *p*_*E*_ and *p*_*I*_ . Orange curves show predicted firing probability when *p*_*I*_ increases linearly with *p*_*E*_ (*p*_*I*_ = *k*_*c*_*p*_*E*_) at different scaling factors *k*_*c*_. Green curves correspond to cases where *p*_*I*_ is held constant. **Inset**, projection of these curves onto the *p*_*net*_ –*p*_*E*_ plane. If each EPSP is suprathreshold, *p*_*net*_ is equivalent to the firing probability *p*_*f*_ , and the plot can be interpreted as the neuron’s input–output relation.

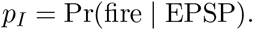

Thus, the joint probability that an EPSP appears in the *I* neuron and that the *I* neuron subsequently fires to generate an IPSP in the *E* cell is *p*_*E*_*p*_*I*_.

Fig. 1A shows four EPSPs (blue) and four IPSPs (red) across seven stimulus trials. For illustrative purposes, EPSPs and IPSPs are assumed to have the same amplitude and latency. An EPSP that coincides with an IPSP is effectively canceled (trials 3 and 6), whereas trials in which the IPSP fails to appear (trials 1 and 4) or appears alone (trials 2 and 7) have no effect. Thus, the probability that an EPSP is not canceled by an IPSP is

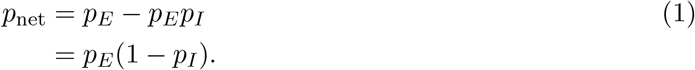

Eq. 1 should be interpreted as a *trial-averaged survival probability*, reflecting both the chance that an excitatory input occurs (*p*_*E*_) and the chance that it is not canceled by coincident inhibition (1 − *p*_*I*_). Over *n* independent trials, the number of uncanceled EPSPs in the reference cell follows a binomial distribution with mean *np*_net_.

The surface plot in Fig. 1A illustrates how *p*_net_ varies with *p*_*E*_ and *p*_*I*_, highlighting key properties of Eq. 1. First, *p*_net_ is nonzero even when *p*_*E*_ = *p*_*I*_, except in the limiting cases *p*_*E*_ = 0 or *p*_*I*_ = 1. Second, *p*_net_ depends on how *p*_*E*_ and *p*_*I*_ co-vary. For example, if *p*_*I*_ scales with *p*_*E*_ as *p*_*I*_ = *k*_*c*_*p*_*E*_ with 0 ≤ *k*_*c*_ ≤ 1, then *p*_net_ increases monotonically with *p*_*E*_ for small *k*_*c*_, but becomes non-monotonic for larger *k*_*c*_ (orange curves). Third, if *p*_*I*_ is fixed, *p*_net_ increases linearly with *p*_*E*_ at a rate given by the slope (1 − *p*_*I*_) (green curves).

The inset shows the orange and green curves projected onto the *p*_net_–*p*_*E*_ plane. If each EPSP is suprathreshold, *p*_net_ corresponds to the firing probability *p*_*f*_ , and the resulting plot represents the I–O relation of the neuron, with *p*_*E*_ serving as a proxy for stimulus intensity. This framework thus links the probabilistic structure of synaptic inputs to the macroscopic I–O behavior of the neuron.

In the following sections, this toy model is extended to more realistic networks with multiple afferents and inhibitory neurons, where cancellation emerges statistically.

### General model

Neuronal responses depend in part on the duration of stimulation. During sustained input, afferents and inhibitory neurons generate sequences of excitatory and inhibitory synaptic inputs, respectively, producing a synaptic barrage in the postsynaptic reference neuron. If the input is sufficiently strong, neurons fire repetitively at a rate determined by the average synaptic current [23]. In contrast, brief stimuli evoke EPSPs and IPSPs that arrive in close temporal proximity, placing the neuron in a regime where spiking is highly sensitive to both the amplitude and timing of synaptic inputs [4]. These two regimes will be treated separately, as the model yields distinct predictions for each. The case with sustained stimulation is examined first.

### Sustained stimuli: oscillatory firing regime

The response of the reference cell to long-duration stimuli was evaluated in three stages (Fig. 2A). First, Poisson-distributed spike trains from *n*_*E*_ external afferents (blue) were generated to produce excitatory synaptic barrages, which were delivered to both the reference cell and the inhibitory (*I*) neurons, modeled as leaky integrate-and-fire (LIF) units. Second, the spike trains evoked in a specified number of *I* neurons (red) were summed to form the inhibitory synaptic barrage. Finally, the excitatory and inhibitory barrages were combined and delivered to the reference cell, and its firing response was measured across a range of conditions.

**Figure 2.**
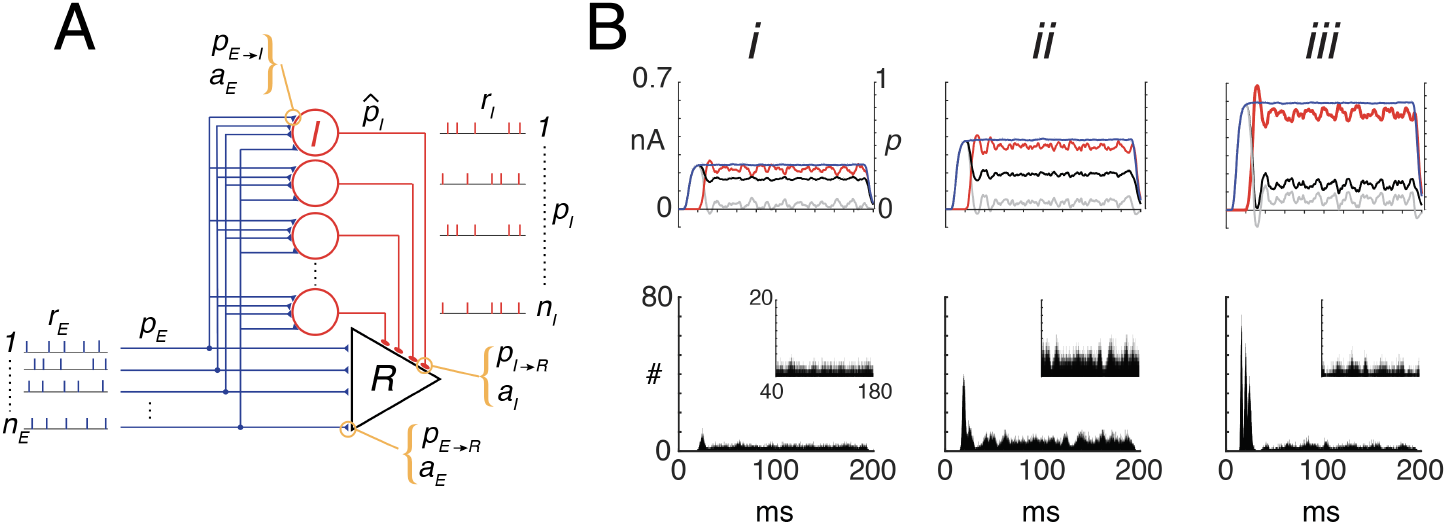
Predicting firing rate in response to sustained stimuli. **A**, Schematic of the network consisting of a reference cell (R, triangle) and *n*_*I*_ inhibitory neurons (red circles). During stimulation, each of the *n*_*E*_ afferent fires with probability *p*_*E*_ at rate *r*_*E*_ for a duration of 0.1 to 1 second. Each afferent spike evoked an EPSC with amplitude *a*_*E*_ in both the reference neuron and each inhibitory neuron with probabilities *p*_*E*→*R*_ and *p*_*E*→*I*_ , respectively. If the excitatory synaptic barrage was sufficiently large, each inhibitory neuron fired with probability 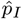 at a rate *r*_*I*_ . The effective probability -which factors in network variables, synaptic strength, and firing rates - that the spikes reach the reference cell is *p*_*I*_ (see text). Finally, the probability that each inhibitory spike evoked an inhibitory postsynaptic current (IPSC) with amplitude *a*_*I*_ in the reference neuron is *p*_*I*→*R*_. In the simulations, *p*_*E*→*R*_ and *p*_*I*→*R*_ are set to 1. **B*i*** , top, Time course of excitatory synaptic current barrage (blue, left ordinate) when *p*_*E*_ (right ordinate) was ramped to a steady-state value of 0.35. The inhibitory barrage (red) plotted as absolute value for comparison) develops after a short delay of 20 ms, eventually reaching a comparable steady-state value. The resulting net current (black), calculated with probability *p*_net_ = *p*_*E*_ (1 − *p*_*I*_), shows a transient peak at stimulus onset followed by a lower steady-state level. In contrast, the unconditioned current obtained by direct subtraction of excitatory and inhibitory currents is nearly zero (gray). **Bottom**, Spike histogram of the reference neuron. **Inset**, magnified view of the tonic firing component. ***ii, iii*** , Same as ***i*** , except with *p*_*E*_ and *p*_*I*_ set to 0.55 and 0.85, respectively. Histograms compiled with bin width=0.01 ms and with 10000 sweeps. **Model parameters**, Simulations used leaky integrate-and-fire neurons (see Methods) with: *n*_*E*_ = 250, *n*_*I*_ = 50, *r*_*E*_ = 50 Hz, *r*_*I*_ ≈ 20–100 Hz. EPSC and IPSC are alpha functions with amplitudes *a*_*E*_ = 10.3, *a*_*I*_ = −10.3 (in pA). The steady-state value of *p*_*I*_ was controlled by fixing the input probability to inhibitory cells (*π*_*E*→*I*_ in Eq. 2) at 0.35 and adjusting the ratio *k*_*n*_ (Eq. 3) to 0.3 (***i***), 0.4 (***ii***), and 0.7 (***iii***). bin width = 0.01 ms

To introduce the key variables manipulated in the simulations, the relationships between the *E* and *I* probabilities and the corresponding synaptic currents are first developed. A detailed description of the model equations and parameters is provided in the Methods and Supporting information; here, only the principal features necessary for interpreting the Results are summarized.

Figure 2A shows the schematic of the feedforward network, with the parameters defined in Table 1.

**Table 1:** Parameters of the feedforward network.

| Symbol | Description |
| --- | --- |
| $p_E$ | Prob. that a stimulus activates an afferent |
| $p_I$ | Effective Prob. that an $I$ cell spike occurs in the reference cell |
| $p_{E \rightarrow R}$ | Prob. that an $E$ synaptic input appears in the reference cell when an afferent fires |
| $p_{E \rightarrow I}$ | Prob. that an $E$ synaptic input appears in an $I$ cell when an afferent fires |
| $p_{I \rightarrow R}$ | Prob. that an $I$ synaptic input appears in the reference cell when an $I$ cell fires |
| $n_E, n_I$ | Numbers of $E$ afferents and $I$ neurons |
| $r_E, r_I$ | Firing rates of $E$ afferents and $I$ neurons |
| $a_E, a_I$ | Amplitudes of $E$ and $I$ postsynaptic currents |
| $q_E, q_I$ | Charge transfers of $E$ and $I$ synaptic events. |

During a prolonged stimulus, each afferent generated a Poisson train of action potentials with mean rate *p*_*E*_ *p*_*E*→*R*_ *r*_*E*_, where *p*_*E*_ is the probability that an afferent becomes active during the stimulus, *p*_*E*→*R*_ is the probability that each spike produces an excitatory postsynaptic current (EPSC) in the reference cell, and *r*_*E*_ is the firing rate conditional on activation. Across afferents or repeated trials, this product represents the effective EPSC rate, and the probability of observing a spike within a time bin of width Δ*t* is approximately *p*_*E*_ *p*_*E*→*R*_ *r*_*E*_ Δ*t*. In this formulation, *p*_*E*_ captures variability across afferents or stimulus presentations, *p*_*E*→*R*_ quantifies synaptic efficacy, and *r*_*E*_ describes the within-trial firing dynamics of an active afferent.

When the *n*_*E*_ afferents fired, each spike evoked an EPSC, whose integral yielded the total charge transfer *q*_*E*_. The mean steady-state excitatory currents to the reference neuron 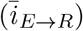 and to the inhibitory neurons 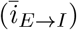 are shown in Fig. 2B (blue; see Sect. 0.2 in the Supporting information). When the afferent input was sufficiently large, the *I* neurons fired at a rate *r*_*I*_ and generated inhibitory synaptic current 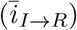 in the reference cell. These mean currents are expressed as

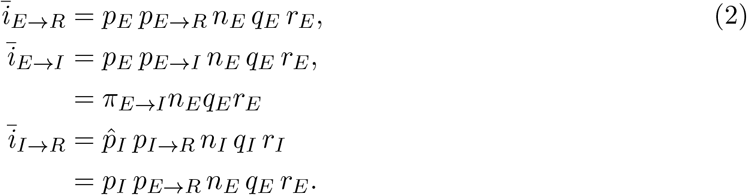

Here, *p*_*E*→*I*_ denotes the probability that an afferent spike evokes an EPSC in an inhibitory neuron, and *p*_*I*→*R*_ denotes the probability that an inhibitory spike evokes an IPSC in the reference cell [47, 48, 49, 8, 50].

The term *p*_*I*_ denotes the *effective* probability that an inhibitory afferent to the reference cell fires during stimulation, or equivalently, the probability that an inhibitory spike reaches the reference cell. It is expressed as a product of factors that relate inhibitory neuron number, synaptic strength, and activity level to those of the excitatory afferents (see Sects. 0 .1 and 0 .2 in the Supporting information):

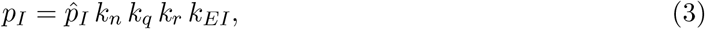

where 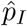 is the probability that an inhibitory neuron fires during the stimulus, 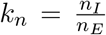 is the inhibitory-to-excitatory input ratio, 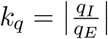 is the ratio of unitary charge transfers, 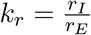is the inhibitory-to-excitatory rate ratio, and 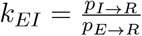 is the synaptic efficacy ratio. By construction, *p*_*I*_ incorporates these relative differences in number, strength, and activity, allowing the inhibitory current (Eq. 2) to be written in terms of *n*_*E*_, *q*_*E*_, *r*_*E*_, and *p*_*E*→*R*_.

The mean net current to the reference cell, under the condition that inhibition is contingent on coincident excitation, is given by (see also Eq. S12 in the Supporting information)

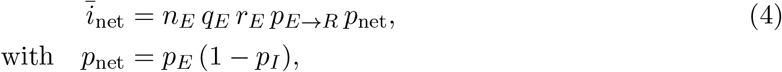

where *p*_net_ denotes the survival probability of excitation—requiring both that an excitatory input occurs with probability *p*_*E*_ and that it is not canceled by coincident inhibition with probability (1−*p*_*I*_) (see Sect. 0.2 in the Supporting information for details). This compact probabilistic form incorporates interneuron recruitment and synaptic reliability, and provides an effective representation of the mean net excitatory drive under feed-forward coupling. In contrast to classical add–subtract models (e.g., for Poisson processes [51]), in which excitation and inhibition are treated as independent and subtracted directly, the present formulation yields a probabilistic measure that, by construction, remains within a normalized range (approximately [0, 1]) for realistic values of identifiable parameters (see Discussion), thereby avoiding the unbounded subtraction of independent terms.

Fig. 2Bi shows the synaptic currents (top) and spike histograms (bottom) recorded from the reference cell during stimulation. Simulations were performed using LIF neurons (see figure caption and Methods for details). The afferent input probability was ramped over time to a steady-state value of *p*_*E*_ = 0.35, producing a barrage of excitatory synaptic currents (blue). Physiologically, this ramp mimics the gradual increase in stimulus intensity before reaching a plateau. When inhibition was modeled as independent of coincident excitation, inhibitory currents (red) were recruited after a short delay and rose to a comparable level. In this case, the unconditioned net current—obtained by direct subtraction of excitatory and inhibitory currents, *i*_*E*_(*t*) − *i*_*I*_(*t*) (gray)—was nearly zero, and no firing occurred (not shown). By contrast, when inhibition was conditioned on coincident excitation, the net synaptic current (black, top) and the firing profile of the reference cell (bottom) displayed a small transient peak followed by a sustained tonic component.

The net current and tonic firing reflected the balance between *p*_*E*_ and *p*_*I*_, in agreement with the toy model (orange curves in Fig. 1B). When both probabilities increased together, 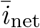 and the tonic firing rate rose to a peak value (Fig. 2B *ii*) before declining (*iii*), and eventually vanished as *p*_*E*_ and *p*_*I*_ approached 1 (see also Fig. 3C).

**Figure 3.**
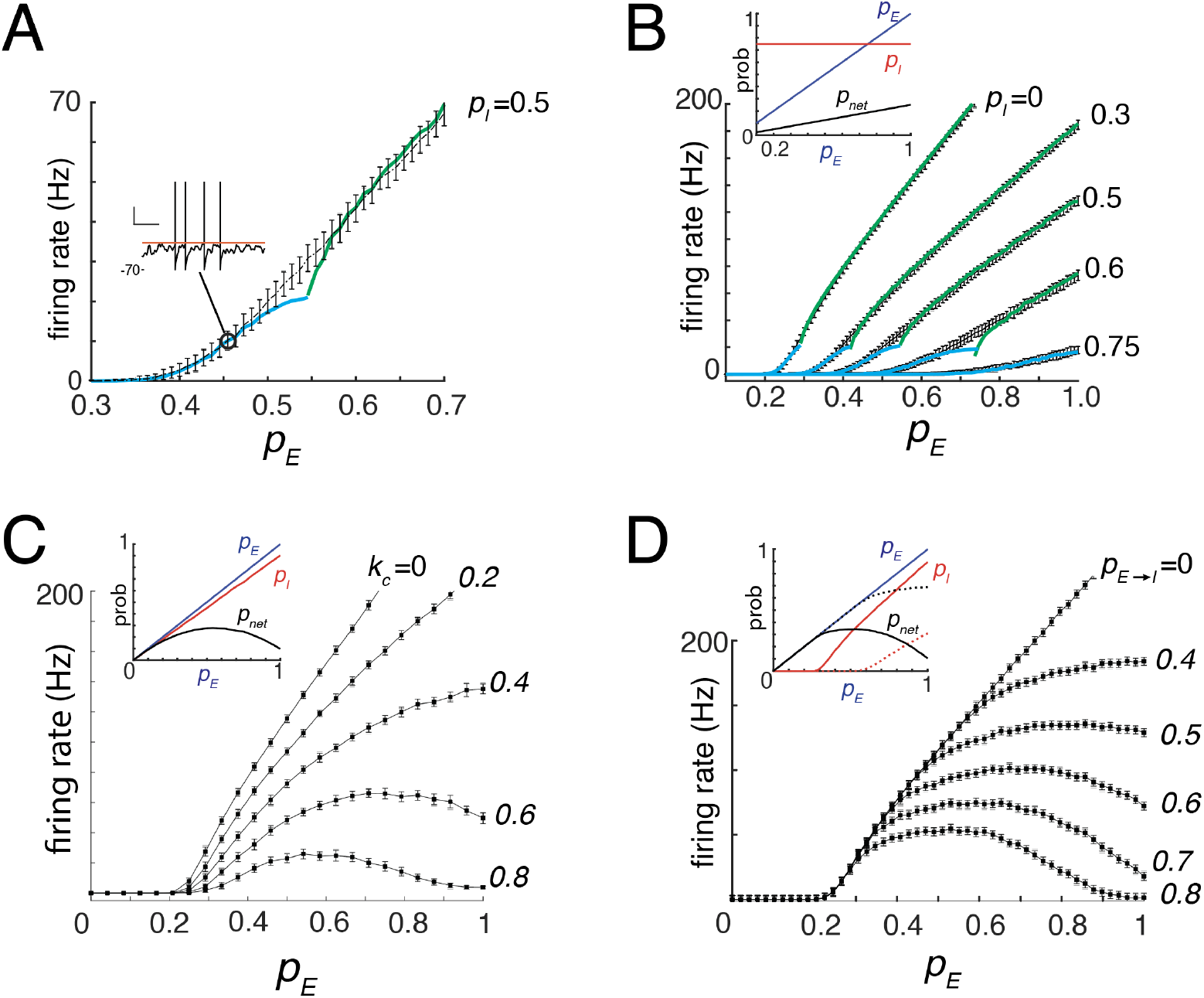
Gain modulation of input–output (I–O) curves. **A**, Representative I–O curve showing mean firing rate (mean ± SD) as a function of *p*_*E*_ , with *p*_*I*_ = 0.46 (*π*_*E*→*I*_ = 0.5, *k*_*n*_ = 0.2, *k*_*r*_ ≈ 2.3). Superimposed are the predicted firing rates in the sub-oscillatory (cyan) and oscillatory (green) regimes. **Inset**, Example membrane potential trace evoked by a sub-rheobase input. Fluctuations in the membrane potential cause threshold crossings (orange line). **B**, I–O curves for different fixed values of *p*_*I*_ . The ratio 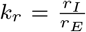 in Eq. 3 was varied by fixing *π*_*E*→*I*_ at specified levels (range: 0–0.8), which in turn modulated *r*_*I*_ . **Inset**, Plots of *p*_*E*_ , *p*_*I*_ (= 0.75), and *p*_net_ versus *p*_*E*_ . **C**, Same as **B**, except *p*_*I*_ increased linearly with *p*_*E*_ , implemented by setting *k*_*n*_ = *k*_*c*_*p*_*E*_ while holding *π*_*E*→*I*_ = 0.4, *k*_*q*_ = 1, and *k*_*r*_ ≈ 1 (see text). **D**, Same as **C**, except *π*_*E*→*I*_ was allowed to vary with *p*_*E*_ for different values of *p*_*E*→*I*_ . **Inset**, *p*_*I*_ remained near zero for low *p*_*E*_ and increased linearly with *p*_*E*_ once threshold was crossed. Solid and dotted curves correspond to *π*_*E*→*I*_ = 0.7 and 0.4, respectively.

Simulations were performed while systematically increasing *p*_*E*_. The resulting input–output (I–O) curve was obtained by plotting the evoked firing rate (mean ± SD) against *p*_*E*_ (Fig. 3A). When the mean net input current 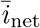 exceeded the rheobase *i*_*rh*_, the reference neuron entered the oscillatory firing regime. In this regime, the average firing rate of the LIF model (green curve) followed the standard analytical solution (see eq. S16 in Supporting information).

Firing could still occur with weak inputs due to voltage fluctuations that occasionally crossed threshold, even when 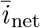 was below rheobase (Fig. 3A, inset). In this fluctuation-dominated (sub-oscillatory) regime (cyan curves), the firing rate was given by

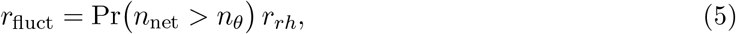

where *r*_*rh*_ is the minimum oscillatory firing rate at rheobase and Pr(*n*_net_ *> n*_*θ*_) is the probability that the excitatory drive exceeds threshold (derivation in sect. 0.2 of the Supporting information). The resulting *r*_fluct_ increased sigmoidally with *p*_*E*_ and approached a maximum near the onset of the oscillatory regime. Across the full range of *p*_*E*_, the overall firing rate was given by the larger of *r*_fluct_ and *r*_osc_.

The model predicts that, for a fixed *p*_*I*_, the slope of the I–O curve scales in proportion to 1 − *p*_*I*_ (Fig. 1B, green; Eq. 1). Consistent with this prediction, increasing *p*_*I*_ (see figure caption for details) caused the I–O curves to exhibit both a reduction in slope and a rightward shift, reflecting a higher activation threshold (Fig. 3B). This condition, in which inhibition was held constant across the full range of *p*_*E*_, is analogous to experiments in which inhibitory neurons are continuously activated optogenetically during sensory stimulation [34]. The slope decrease corresponds to multiplicative gain modulation, whereas the threshold shift reflects an additive effect of persistent inhibition, requiring stronger excitation to elicit firing. These effects were observed in both the oscillatory and fluctuation-dominated regimes and were captured by the analytical expressions for *r*_osc_ (green) and *r*_fluct_ (cyan). Together, these results demonstrate that inhibition modulates the I–O relationship through a combination of multiplicative and additive mechanisms [13, 29].

The I–O curves exhibited either monotonic or non-monotonic increases with *p*_*E*_, depending on how strongly inhibition co-varied with excitation (Fig. 3C), consistent with model predictions (Fig. 1B, orange curves). Co-variation of *p*_*I*_ with *p*_*E*_ was implemented by allowing *k*_*n*_ to increase linearly with *p*_*E*_ (*k*_*n*_ = *k*_*c*_*p*_*E*_, *k*_*c*_ ∈ [0, 1]) (inset). Physiologically, this corresponds to the progressive recruitment of inhibitory neurons with increasing stimulus intensity. For small *k*_*c*_, firing rates increased monotonically with *p*_*E*_ (Fig. 3C), whereas as *k*_*c*_ approached 1, the I–O curves flattened and eventually became non-monotonic. Notably, the minimum *p*_*E*_ required to evoke firing (∼ 0.2) remained unchanged, while the slope of the rising phase decreased. Thus, when *p*_*E*_ was restricted to the range in which the I–O curve increased, gain modulation was effectively multiplicative.

Finally, the model predicts changes in the I–O curves of the reference cell under more physiological conditions, in which inhibition strengthens as excitation increases. In earlier simulations, *p*_*I*_ was manipulated by fixing the drive to inhibitory neurons (Eq. 2). Here, the effective drive *π*_*E*→*I*_ was allowed to increase with *p*_*E*_, mimicking increased drive to inhibitory neurons with rising stimulus intensity (Fig. 3D, inset). The slope of this recruitment was controlled by varying *p*_*E*→*I*_, which physiologically corresponds to changes in the efficacy of the afferent synapse onto inhibitory neurons. Increasing *p*_*E*→*I*_ shifted the *p*_*I*_ curve leftward (dotted to solid red, inset), causing the I–O curves of the reference neuron to become progressively more non-monotonic (Fig. 3D). The curves shared a common threshold and overlapped at low *p*_*E*_ before diverging at higher values. Although these changes do not conform to classical forms of gain modulation, they can still generate multiplicative effects on tuned inputs (see below).

### Correction for conductance effects

A key assumption in Eqs. 2, and 4 is the linear summation of excitatory and inhibitory inputs. This assumption is violated when synaptic inputs alter the total membrane conductance, since the net current at rest can differ substantially from that during depolarized states, leading to errors in predicted firing rates. To account for this effect, a conductance-dependent adjustment was derived (Sect. 0.2 in the Supporting information). Incorporating this correction substantially improved prediction accuracy (Fig. S2) while preserving the probabilistic formulation, allowing the same framework to be applied under conductance-based synapses.

### Gain modulation of tuned responses

The slope changes in the I–O curves described above (Figs. 3B–D) suggest mechanisms for multiplicative gain modulation of neural responses [27, 28, 52]. To test this, simulations were performed with *p*_*E*_ following a Gaussian profile representing tuned sensory input (Fig. 4A). Three conditions were examined: (mode 1) fixed *p*_*I*_; (mode 2) *p*_*I*_ increasing linearly with *p*_*E*_; and (mode 3) *p*_*I*_ following the I–O curve of the inhibitory neurons. In all cases, the peak input (*p*_*E*_ = 0.35) produced modulated tuning curves with peak firing rates within 50% of the control (60 Hz), consistent with experimental data [27, 28, 52].

**Figure 4.**
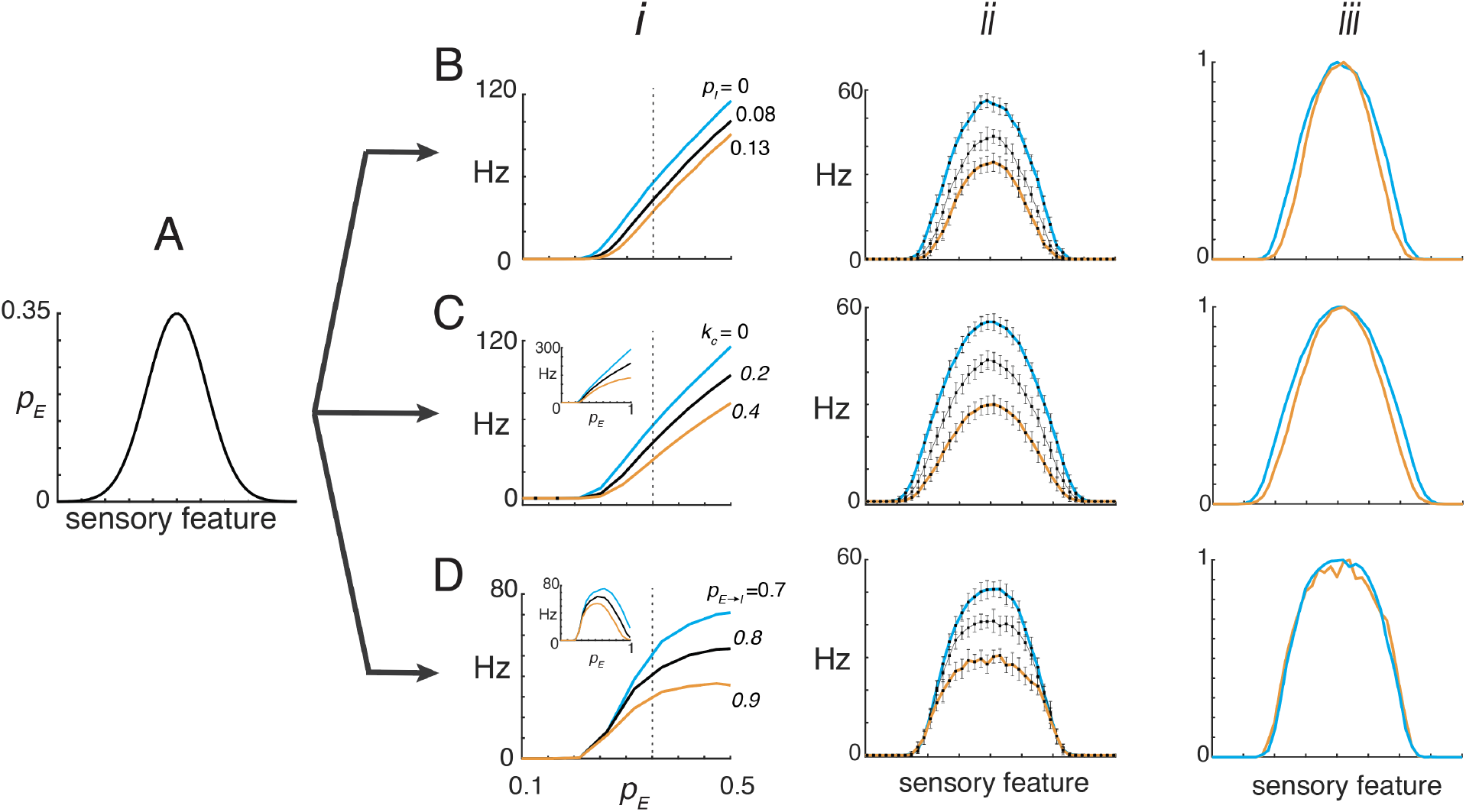
Gain modulation of tuned responses. **A**, Simulations in which *p*_*E*_ followed a Gaussian profile, representing a tuned sensory input. **B**, Case with constant *p*_*I*_ set to 0, 0.08, or 0.13. ***i*** , I–O curves used to transform the input. ***ii*** , Resulting tuned responses of the reference neuron. ***iii*** , Tuned responses normalized to a peak value of 1. **C**, Same as in **B**, except *p*_*I*_ increased linearly with *p*_*E*_ . This was implemented by setting *k*_*n*_ = *k*_*c*_ *p*_*E*_ with *k*_*c*_ = 0, 0.2, or 0.4 at a fixed *π*_*E*→*I*_ . **Inset**, Corresponding I–O curves across the full range of *p*_*E*_ . **D**, Same as in **C**, except *π*_*E*→*I*_ varied with *p*_*E*_ at different rates by setting *p*_*E*→*I*_ = 0.7, 0.8, or 0.9.

For mode 1, relatively small values of *p*_*I*_ were sufficient to produce multiplicative gain modulation. Although the I-O curves showed both slope and threshold changes (Fig. 3B), the small range of *p*_*I*_ that was used primarily affected the slope (Fig. 4B, left), resulting in a reduction in tuning curve amplitude (middle) with minimal change in width (right). Physiologically, such low *p*_*I*_ values (eq. 3) could arise from a small number of inhibitory neurons relative to afferents (low *k*_*n*_), weak synaptic inhibition (low *k*_*q*_), and/or low inhibitory firing rates (low *k*_*r*_).

In mode 2, multiplicative gain modulation was achieved by setting the scalar *k*_*c*_ in the relationship *p*_*I*_ = *k*_*c*_*p*_*E*_ to modest values. The resulting I–O curves were monotonic, with progressively reduced slopes as *k*_*c*_ increased (Fig. 4C, left and inset). The peaks of the corresponding tuned responses (middle) varied with *k*_*c*_ and, when normalized, superimposed (right). To achieve a linear relationship between *p*_*I*_ and *p*_*E*_ under physiological conditions would require a fixed *π*_*E*→*I*_ across input intensities and low inhibitory firing thresholds to ensure engagement even for small *p*_*E*_ values.

In mode 3, multiplicative gain modulation occurred but only within a limited range of *p*_*E*_. At low *p*_*E*_, the I–O curves overlapped substantially (Fig. 4D, left), and divergence required strong inhibition at moderate *p*_*E*_, where slope differences emerged. Under these conditions, the I–O curves became non-monotonic (inset). When *p*_*E*_ exceeded the range of the rising phase, firing shifted to the decaying portion of the curve, producing a central dip and bimodal tuning (Fig. S3). When restricted to the rising phase, however, peak responses could be modulated (middle) with minimal changes in tuning width (right).

### Temporal firing profiles

Neuronal firing patterns can encode distinct stimulus features or task-related components [36, 53, 54]. As the stimulus changes, the drive to the neuron—reflected in variations of *p*_*E*_—also changes. To examine how such temporal profiles depend on E–I interactions, it is necessary to systematically vary the time-dependent inputs. Controlling the inhibitory drive is challenging, however, because its onset and magnitude depend on how inhibitory neurons are recruited by the stimulus (Fig. 2B). This recruitment, in turn, depends on the biophysical properties of inhibitory neurons [55] and on the amplitude of their excitatory inputs [56, 57, 58]. The firing profile of the reference cell also depends on the relative timing of EPSPs and IPSPs: in most cases, IPSPs lag EPSPs [2, 18, 40], although they can also precede them under certain conditions [59, 60].

Replicating the full range of possible time-varying relationships between *p*_*E*_(*t*) and *p*_*I*_(*t*) would require detailed modeling of inhibitory neuron dynamics, which is beyond the scope of this study. To allow direct and independent control of *p*_*I*_(*t*), the inhibitory neurons were bypassed, and inhibitory inputs were generated in the same way as the excitatory afferents. The excitatory input, together with the conditioned inhibition *p*_*E*_(*t*)*p*_*I*_(*t*), was then delivered to the reference cell. This abstraction isolates the temporal interaction between excitation and inhibition, enabling their overlap to be examined in a controlled and systematic manner.

Both *p*_*E*_(*t*) and *p*_*I*_(*t*) were ramped up to the same steady-state value and then ramped down (blue and red dashed curves in Fig. 5A–C, bottom panels). The simulation parameters were identical (see Fig.6 captions and Methods for details) except that the relative onsets were varied. The barrages were calculated as above and delivered to the LIF neuron, and firing histograms were compiled across repeated trials (top panels).

**Figure 5.**
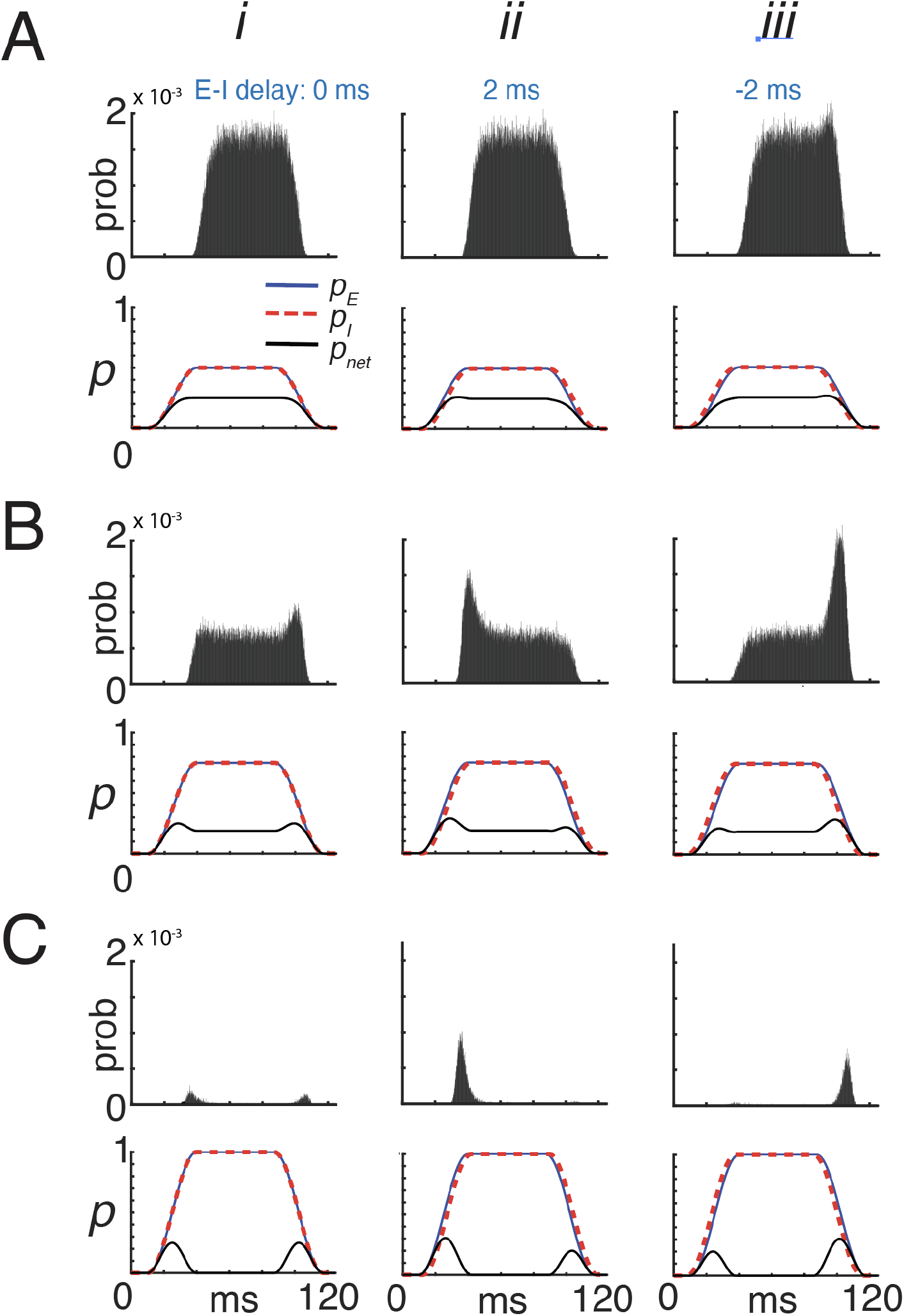
Effects of inhibitory magnitude and timing on temporal firing. **A*i*** , *bottom*: Excitatory and inhibitory input probabilities, *p*_*E*_ (*t*) (blue) and *p*_*I*_ (*t*) (red), rose and fell together with no delay, each reaching a steady-state value of 0.5. The computed *p*_net_(*t*) = *p*_*E*_ (*t*) [1 − *p*_*I*_ (*t*)] is superimposed (black). *Top*: the corresponding spike histogram shows tonic firing in the reference neuron. ***ii–iii*** , Same as ***i*** , except *p*_*I*_ (*t*) lagged or led *p*_*E*_ (*t*) by 2 ms, respectively. **B**, Same as in **A**, but with larger steady-state probabilities (0.75). **C**, Same as in **A**, but with steady-state probabilities equal to 1. **Model parameters:** LIF neuron as in Methods. Per-afferent input intensities were *λ*_*E*→*R*_(*t*) = *p*_*E*_ (*t*) *r*_*E*_ and *λ*_*I*→*R*_(*t*) = *p*_*E*_ (*t*) *p*_*I*_ (*t*) *r*_*E*_ , with *r*_*E*_ = 50 Hz and population sizes *n*_*E*_ = *n*_*I*_ = 250, bin width = 0.01 ms.

A wide range of temporal firing patterns was generated by varying the magnitudes and relative timing of *p*_*E*_(*t*) and *p*_*I*_(*t*). In these simulations, the time courses of *p*_*E*_(*t*) and *p*_*I*_(*t*) were identical (Fig. 5A–C, bottom panels) but differed in their temporal lags (*i–iii*). For moderate inputs with no lag (A, *i*), the computed *p*_net_(*t*) (black) closely followed the time courses of *p*_*E*_(*t*) (blue) and *p*_*I*_(*t*) (red dashed), ramping to a steady level before decaying. The reference neuron responded with delayed tonic firing (top).

With larger input magnitudes (B), *p*_net_(*t*) exhibited transient peaks at stimulus onset and offset, accompanied by a reduced steady-state level. Further increases in steady-state amplitude (C) eliminated the tonic component entirely, leaving only onset and offset peaks, since *p*_*E*_(*t*)[1−*p*_*I*_(*t*)]*/*= 0 only during those intervals. Accordingly, the reference neuron fired exclusively at stimulus onset and offset (top).

When *p*_*I*_(*t*) lagged *p*_*E*_(*t*) by 2 ms (*ii*), similar results were obtained, except that the onset peak of *p*_net_(*t*) was larger than the offset peak (Fig. 5C). This produced pronounced firing at stimulus onset—greater than that observed in the no-delay condition (*i*)—and no firing at stimulus offset. Conversely, when *p*_*I*_(*t*) preceded *p*_*E*_(*t*) (*iii*), spiking occurred only at stimulus offset.

### Brief Stimuli: Transient firing regime

Brief stimuli, such as tone pips in the auditory system [59, 2, 33], light flashes in the visual system [61], or whisker deflections in the somatosensory system [6, 4], evoke a volley of compound EPSPs, followed by compound IPSPs from inhibitory neurons after a small delay. Whether the postsynaptic cell fires depends both on the relative magnitude and timing of these inputs.

To examine the effects of *E*–*I* balance and timing on firing probability, simulations were performed in the same network as above. A brief stimulus evoked EPSPs in both the reference and *I* neurons whose arrival times followed a Gaussian distribution (Fig. 6A*i*, top panel, blue histogram).

**Figure 6.**
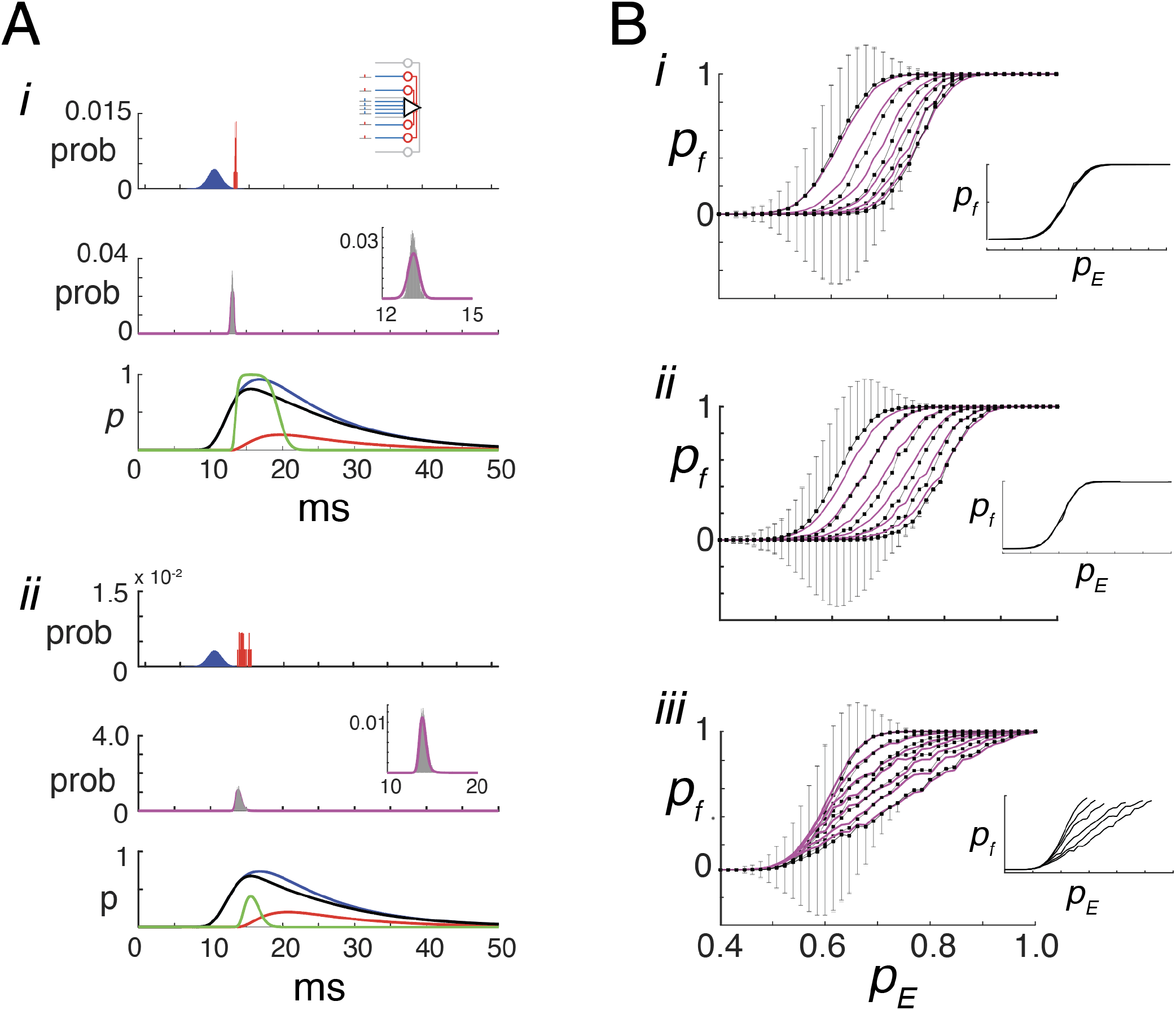
Predicting firing probability for transient stimuli. **A*i***,**top**, probability distributions of arrival times of EPSP (blue) and IPSP (red) in the reference cell. **Middle**, predicted firing probability (magenta) overlaid on the spike time histogram (gray); **Inset**, magnified view. **Bottom**, superimposed traces of 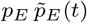 (blue), 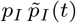 (red), net input 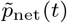 (black), and threshold-crossing probability (green). *p*_*E*_ = 0.95, *k*_*n*_ = 0.2, *k*_*a*_ = 1, *k*_*EI*_ = 1. ***ii*** , same but with *p*_*E*_ = 0.75 . **B**,***i*** , I-O curves showing mean firing probability (±SD) vs. *p*_*E*_ , for fixed *p*_*I*_ values (0 ≤ *p*_*I*_ ≤ 0.5). The *p*_*I*_ level was specified by adjusting and then fixing *k*_*n*_ ∈ [0, 0.5]. Other parameters are kept constant: *p*_*E*→*R*_ = 1, *p*_*E*→*I*_ = 0.8, *a*_*E*_ = *a*_*I*_ = *±*10.3 pA, which produced PSPs with amplitude *±*250*µV* . Predicted values (magenta) are overlaid. Increasing *p*_*I*_ shifts the curve rightward without changing slope. **Inset**, I-O curves aligned by threshold. ***ii*** , same but with *p*_*I*_ increasing linearly with *p*_*E*_ , implemented by setting *k*_*n*_ = *k*_*c*_*p*_*E*_; *k*_*c*_ ∈ [0, 1] while holding *π*_*E*→*I*_ =. ***iii*** , same but with *p*_*E*→*I*_ = *k*_*p*_*p*_*E*_; 0.85 ≤ *k*_*p*_ ≤ 1, *k*_*n*_ = 1. **Model parameters:** *n*_*E*_ = 100, *n*_*I*_ = *k*_*n*_ *n*_*E*_ . Histograms compiled over 5000 trials with Bin width = 0.01 ms

When the *I* neurons fired, they generated IPSPs in the reference cell a short time later, with a narrower temporal distribution (red). In response to these EPSPs and IPSPs, the reference cell fired action potentials with a narrower distribution (middle, gray) than the input, consistent with experimental observations [4].

The instantaneous excitatory and inhibitory probabilities (blue and red traces in Fig. 6A, bottom panel) were computed from the compound EPSPs and IPSPs using convolution expressions derived in Supporting Information 0.2. These are given by

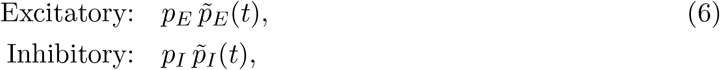

where 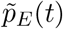 and 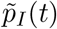were obtained by convolving peak-normalized unitary PSPs with the arrivaltime histograms of afferent and inhibitory spikes. The effective probability *p*_*I*_ is as defined above, except that the ratio of inhibitory to excitatory amplitudes 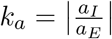 replaces the ratio of charges *k*_*q*_. The time-varying excitatory (blue) and inhibitory (red) probabilities thus follow the same temporal profiles as the compound EPSP and IPSP, respectively (Fig. 6A*i*, bottom panel).

The instantaneous probability density of net excitation—that is, the component of the excitatory drive that is not canceled by inhibition (black curve in Fig. 6A, bottom panel)—is given by

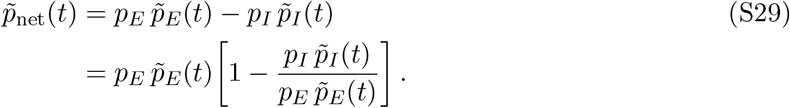

The ratio 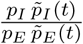 quantifies the instantaneous balance between inhibition and excitation. This time-dependent formulation is directly analogous to the expression for sustained stimulation, where *p*_net_ = *p*_*E*_(1 − *p*_*I*_) represents the steady-state probability that excitation survives inhibition. The expression remains well defined even when 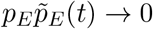, because in a feedforward circuit inhibition co-varies with excitation and both vanish in the absence of a stimulus (i.e., when *p*_*E*_ = 0). To preserve its interpretation as a probability, the constants defining *p*_*I*_ should be chosen such that 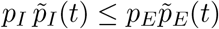 for all *t*; alternatively, a lower bound of zero can be imposed.

The timing of action potentials is computed in two stages (see sect. 0.2 in Supporting information). First, the probability that the net excitatory input at a given time exceeds the threshold required for firing was computed from *p*_*net*_(*t*) (bottom, green curve; eq. S30). Second, this probability was used to derive the expected firing time (Eq. S31), accounting for the constraint that a neuron can fire only once and only if it has not already fired. This procedure yields the predicted firing probability (middle panel, magenta curve), which closely matches the spike histogram (gray, inset). Similar results were obtained for a weaker stimulus (Fig.6A,ii) and when the synaptic inputs were delivered as conductances (Fig. S4 in Supporting information).

Modulating inhibitory strength led to changes in the I–O curves that deviated from model predictions. The total probability of firing *p*_*f*_ was calculated either from the area under the spiking histogram (gray) or from the predicted probability (magenta). Figure 6*i* shows *p*_*f*_ plotted against *p*_*E*_ for different levels of *p*_*I*_, analogous to the green curves in Fig. 1B. The value of *p*_*I*_ was controlled by adjusting and then fixing *k*_*n*_ in Eq. 3 so that *p*_*I*_ remained constant across values of *p*_*E*_ . As *p*_*I*_ increased, the level of *p*_*E*_ required to evoke firing also increased, resulting in a rightward shift of the I–O curves without a change in slope. When aligned by their respective thresholds, the curves collapsed onto a single profile (inset). Thus, unlike in the toy model or under sustained input conditions, the effect on the I–O relationship was purely additive.

Similar results were obtained when *p*_*I*_ increased linearly with *p*_*E*_ (as in Fig.3C). This was implemented by letting *k*_*n*_ increase as: *k*_*n*_ = *k*_*c*_*p*_*E*_. As the slope of *p*_*I*_ vs *p*_*E*_ increased, the curves shifted to the right (*ii*), but unlike the toy model (Fig. 1B, orange curves) and tonic input (Fig. 3C) , remained monotonic with no change in slope (inset). This occurred because the *I* did not cancel the *E* completely: a time window always existed between the onsets of 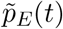 and 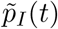 during which action potentials could be evoked. Thus, the effect on the I-O curve is also additive.

In Fig. 6B*(iii)*, the inhibitory input probability *p*_*I*_ was modulated by varying *p*_*E*→*I*_ so that, as in the sustained-stimulus condition (Fig. 3D, inset), the resulting *p*_*I*_ reflected the I–O function of the inhibitory (*I*) neurons. As *p*_*E*→*I*_ increased, the slopes of the I–O curves of the reference cell decreased, whereas their thresholds remained constant. Unlike in the simple model, the curves remained monotonically increasing, with those obtained under weaker inhibition saturating at high *p*_*E*_. Thus, under these conditions, the effect on the I–O curve was predominantly multiplicative within the range preceding saturation (inset).

## Discussion

This study aimed to elucidate the principles governing stimulus-driven excitatory–inhibitory (E–I) interactions in feed-forward inhibitory circuits. By combining excitatory and inhibitory effects within a probabilistic framework, a simple relationship was derived that predicts how E–I balance shifts during stimulation. This formulation links those shifts to both cellular and network-level variables, offering mechanistic insight into how inhibition shapes neuronal I–O functions, modulates gain, and influences overall response dynamics.

A probabilistic formulation offers several theoretical advantages over classical approaches that rely on summing synaptic currents to predict firing [51]. First, the multiplicative expression introduced here captures the survival of excitatory inputs under feed-forward inhibition in a form that is both biologically interpretable and analytically tractable. Because probabilities are inherently bounded between 0 and 1, the resulting I–O curves are automatically rectified and sigmoidal—features that are often imposed manually in models to prevent negative or unbounded firing probabilities or rates [10, 19, 30, 62, 63, 64].

Second, the net probability *p*_net_ can be used directly in analytic expressions to derive I–O relationships and characterize gain control. The reduced form naturally incorporates the variables that determine *p*_*I*_ (such as interneuron recruitment and synaptic reliability) and automatically scales with *p*_*E*_, eliminating the need for manual adjustment when analyzing I–O curves. Because *p*_net_ closely tracks temporal firing patterns, it can be used directly, or as input to Poisson point-process models [65], to efficiently predict stimulus-evoked responses with realistic temporal statistics.

Third, this framework explicitly incorporates synaptic noise, yielding smoothly rising I–O curves without additional assumptions. The sources of variability are well defined and can, in principle, be used to compute statistical properties of both inputs and outputs, linking fluctuations in synaptic drive to the variability and reliability observed in neuronal firing.

### Relation to previous work

The model’s I–O curves exhibit two distinct regimes. For weak stimuli (low *p*_*E*_), firing occurs in a sub-oscillatory regime driven primarily by input variability, producing a sigmoidal rise. This behavior aligns with observations in visual cortex, where input summation is supralinear for weak stimuli and sublinear for strong ones [64, 66]. The underlying mechanisms may differ, however, since the cortical responses reflect recurrent connectivity, which is absent in the present feed-forward model. As input strength increases, the system transitions to an oscillatory regime, in which the I–O curve may continue to rise, flatten, or become non-monotonic. The specific shape of the curve is determined by how inhibition scales with excitation, consistent with previous modeling results [19].

The model highlights key variables that govern the transformation of synaptic input into spiking output. Understanding this transformation has practical implications, as it may enable the inference of intracellular dynamics from extracellular recordings [67]. Although direct inversion of the model equations is not feasible due to the number of interacting variables (e.g., *p*_*E*_, *p*_*I*_, *k*_*n*_, *k*_*q*_, *k*_*r*_), the derived relationships can nevertheless inform which parameters must be independently measured or constrained to obtain reliable estimates.

Multiplicative gain modulation of I–O curves is essential for maintaining tuning acuity across different cognitive or behavioral states [27, 28, 68]. The present model reveals multiple mechanisms for achieving such modulation by varying how *p*_*I*_ scales with *p*_*E*_ (Fig. 3B–D). Unlike previous models [10, 30, 13], these effects arise without invoking recurrent connectivity, synaptic noise, or conductance-based mechanisms. Moreover, the framework provides a means to predict neuromodulatory influences based on their actions on synaptic or biophysical parameters [69, 70].

For brief stimuli, in which firing is strongly influenced by the delay between excitation and inhibition [4, 7, 59], multiplicative gain emerged only when the drive to inhibitory neurons increased with *p*_*E*_ (Fig. 6B*iii*)—a condition likely to occur with natural stimuli. Optogenetic activation of inhibitory neurons during brief sensory stimulation [33] most closely resembles the simulations with fixed *p*_*I*_ (Fig. 6B*i*), which predict additive modulation of the I–O curves. The mixed multiplicative and additive effects observed experimentally may therefore reflect the extent to which optogenetic activation overrides the normal physiological recruitment of inhibitory neurons. For prolonged stimuli, the model predicts that modulation is predominantly multiplicative, consistent with observations during sustained optogenetic activation of inhibitory neurons [34].

The model reproduced many of the temporal firing patterns observed in cortical and subcortical regions [35, 36, 37, 38, 39, 46, 53] by adjusting the magnitude and timing of the excitatory and inhibitory probabilities (Fig. 5). A key prediction is that transient firing and non-monotonic I–O curves observed experimentally [16, 35] need not arise from disproportionately strong inhibition. Both phenomena can emerge even when *p*_*E*_ and *p*_*I*_ increase in a balanced or proportional manner (Figs. 2B, 3C–D, 5). The model further predicts that onset–offset responses can occur when both excitatory and inhibitory probabilities approach unity, with small shifts in inhibitory delay producing distinct temporal response components (Fig. 5; [42]). Whether similar mechanisms operate in cortex remains unclear, as onset and offset responses may be inherited from subcortical pathways [53, 40] or generated *de novo* within cortical circuits [22, 41, 44].

### Limitations

The theory relies on several key assumptions. First, the calculation of *p*_net_ assumes that inhibitory inputs sum linearly with, and effectively cancel, excitatory inputs. Although the sublinear summation associated with conductance changes can be minimized through a correction term, the model does not account for heterogeneity in the amplitudes of individual EPSPs and IPSPs. For analytical simplicity, IPSP amplitudes were expressed as fixed fractions of the corresponding EPSPs. In reality, postsynaptic potential size depends on several factors, including synaptic location and the identity of the presynaptic population. For example, thalamocortical synapses are typically stronger than cortico-cortical synapses [71]. Moreover, IPSPs at the soma can, in principle, shunt or cancel multiple small EPSPs originating in distal dendrites.

Second, the constants (*k*_*n*_, *k*_*EI*_, *k*_*q*_, *k*_*r*_) that scale *p*_*I*_ were chosen (Table 2) so that their product lies between 0 and 1, ensuring that *p*_*I*_ remains interpretable as a probability. Although precise values for these constants are unlikely to be fully known in any given system, reasonable estimates can be drawn from the literature (*k*_*n*_, [72, 73]; *k*_*a*_, *k*_*q*_, [8, 9, 74]; *k*_*r*_, [8, 9, 74]; *k*_*EI*_, [49, 57, 75]). In addition, *k*_*q*_ or *k*_*a*_ may vary dynamically, reflecting short-term plasticity of excitatory and inhibitory synapses [9, 55]. For modeling purposes, parameters should be constrained so that the product remains below unity. As an additional safeguard, a smooth saturating transform, such as *p*_*I*_(*t*) = 1 − exp[−*k*_*n*_*k*_*EI*_*k*_*q*_*k*_*r*_ *p*_*I*_(*t*)], can be applied to preserve the probabilistic interpretation of *p*_*I*_ even when the product exceeds one.

**Table 2:**
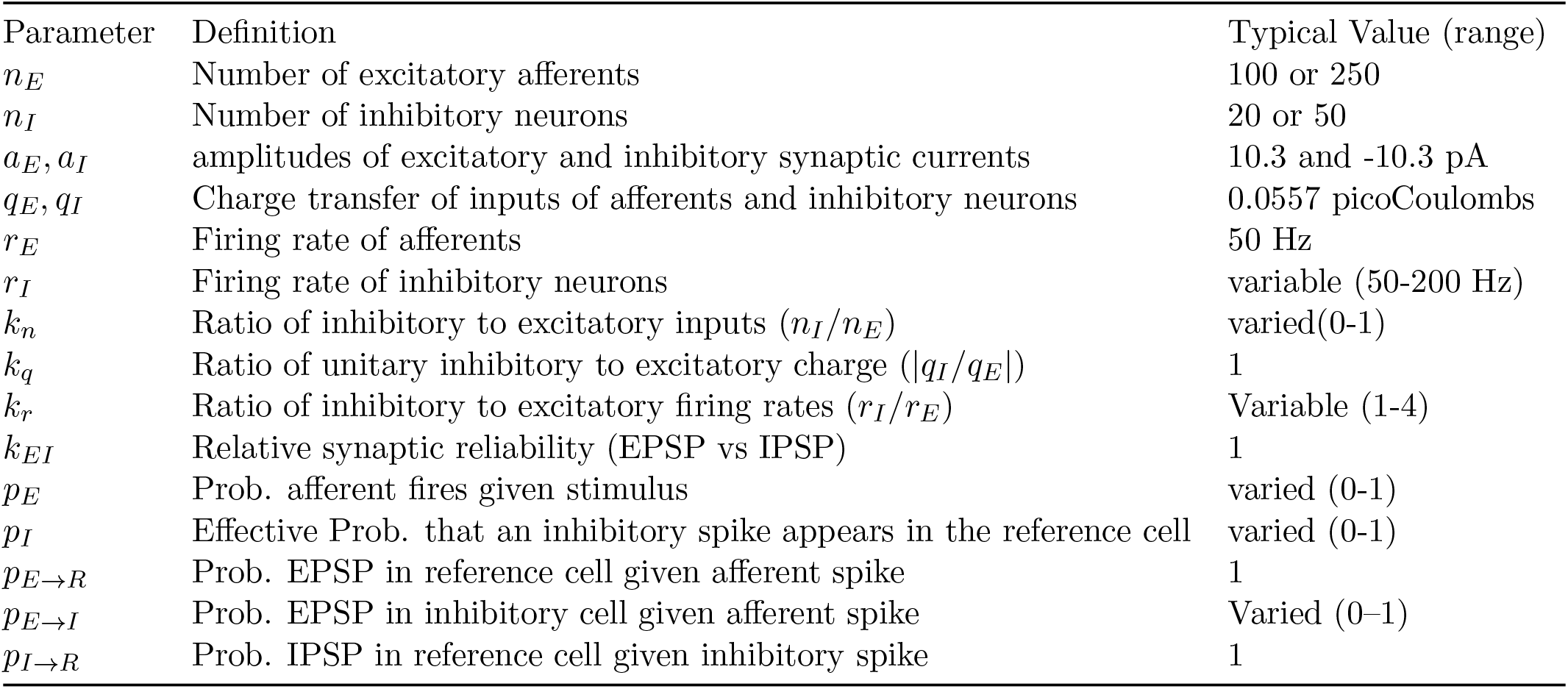
Typical network parameter values used in simulations.

Third, the model assumes that recurrent connections do not substantially contribute to the evoked firing rate. This assumption is likely valid for brief stimuli, during which recurrent activity has limited influence on firing [76]. In the cortex, the connection probability between excitatory neurons is relatively low [9, 55]. Consequently, if the active region is spatially restricted—as in tuned responses driven by topographically organized inputs—the small number of recruited local excitatory neurons would exert minimal impact, consistent with simulation results [76]. Nonetheless, incorporating recurrent connectivity will be important for extending the model to more general network configurations.

Fourth, spontaneous activity—whether intrinsically generated or originating from neurons outside the feedforward pathway—is not explicitly included in the present analysis. Background activity can alter the initial conditions of the network, influence transient responses [77, 78], and contribute to steady-state firing. In principle, such effects could be incorporated into the probabilistic framework if expressed in terms of effective input probabilities, although this would require additional theoretical development.

Finally, a potential limitation of the framework is that spiking is described as being determined solely by inputs within the current time bin, without explicit dependence on prior activity. This simplification is partly mitigated by the model’s structure. For brief stimuli, the probability of a spike at a given time is defined conditionally on no spikes having occurred in earlier bins, thereby incorporating past spiking history implicitly into the first-spike probability. In addition, the time-dependent changes in probability follow the shape of the underlying synaptic potentials, which reflect recent input dynamics. For long-duration stimuli, the analysis focuses on the average synaptic current, which depends primarily on the overall rate and distribution of synaptic events. In this regime, the precise temporal sequence of individual inputs becomes less critical, as the mean current is determined by their statistical properties.

Despite these limitations, the model effectively captures many key aspects of stimulus-evoked responses and can serve as a foundation for developing formal mathematical analyses [79] to study more complex network configurations.

## Methods

### Network parameters

The circuit consists of a reference neuron, whose firing activity serves as the model’s output, and 20-50 local inhibitory neurons. Both cell types received *n*_*E*_ afferent from an excitatory source, with the reference cells also receiving inputs from *n*_*I*_ interneurons. Typical values of the variables used in the simulations are listed in table 2.

### Leaky integrate-and-fire parameters

Simulations were performed in the Matlab programming environment. Neurons were modeled as standard leaky integrate-and-fire (LIF) units governed by

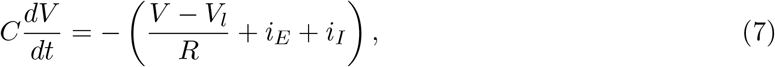

where *R* = 75 MΩ, *V*_*l*_ = −70 mV is the resting potential, *τ*_*m*_ = 10 ms is the membrane time constant, *C* = 133 pF is the capacitance, and the action potential threshold was set to −55 mV. After an action potential, the membrane potential was reset to *V*_*l*_. The terms *i*_*E*_ and *i*_*I*_ denote the total excitatory and inhibitory synaptic currents, respectively (in nA). The integration time step (bin width) was Δ*t* = 0.01 ms.

Each unitary synaptic current *u*_*x*_(*t*), evoked by a single presynaptic spike, was described by a scaled alpha function:

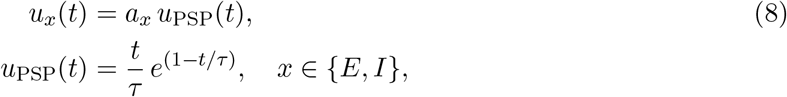

where *τ* = 2 ms and the peak amplitude was normalized to 1. For current-based simulations, *a*_*E*_ = 10.3 pA and *a*_*I*_ = −10.3 pA. For conductance-based simulations, *a*_*E*_ = 0.147 nS and *a*_*I*_ = 1.045 nS, with reversal potentials *V*_*E*_ = 0 mV and *V*_*I*_ = −80 mV. Under these conditions, the corresponding excitatory and inhibitory postsynaptic potentials (PSPs) had amplitudes of approximately +250 *µ*V and −250 *µ*V, respectively, when measured at the resting potential.

### Stimulus parameters

#### Sustained stimulus

For sustained stimulation, excitatory synaptic barrages to the reference and inhibitory neurons were generated by creating a spike train according to a Poisson process with total rate *λ* = *n*_*E*_*p*_*E*_*r*_*E*_ where *r*_*E*_ = 50 Hz is the average firing rate of a single afferent and *n*_*E*_ = 250 is the number of afferents. The excitatory barrage to the inhibitory cells was adjusted by scaling (multiplying) the total rate by *p*_*E*→*I*_. The excitatory barrage (0.1–1 s duration) was then obtained by convolving the spike trains with *u*_*E*_.

To construct the inhibitory barrage, the excitatory barrages were delivered to 3*n*_*I*_(= 3*k*_*n*_*n*_*E*_) inhibitory cells, and the resulting spike trains were stored in a matrix. From this set, *p*_*E*_*n*_*I*_ trains were randomly selected, summed, and subsequently convolved with *u*_*I*_. This inhibitory barrage was then delivered together with the excitatory barrage to the reference neuron. The average firing rate and peristimulus time histograms of the reference neuron were computed over 100–10000 trials, with each trial using different realizations of the synaptic barrages.

To examine gain modulation of receptive fields (Fig. 4), the excitatory input probability was modeled as a Gaussian function,

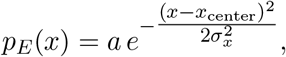

where *p*_*E*_(*x*) denotes the *effective* excitation probability at stimulus feature *x, x*_center_ is the preferred stimulus, and *σ*_*x*_ is the tuning width. This spatially tuned *p*_*E*_(*x*) was then used to generate the excitatory drive to both the reference cell and the inhibitory neurons, following the procedures described above. Average excitatory and inhibitory synaptic currents, together with the evoked firing rate, were computed over 100 trials and plotted as functions of *x*.

In a subset of simulations (Fig. 5), the inhibitory population was bypassed. In these cases, the inhibitory input was generated using the same procedure as for the excitatory input and, after appropriate scaling, was delivered directly to the reference neuron. Both excitatory and inhibitory spike trains were modeled as inhomogeneous Poisson processes with time-varying probabilities, producing instantaneous rates

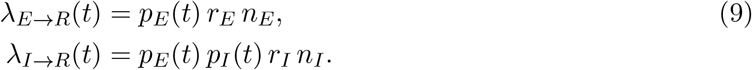

The functions *p*_*E*_(*t*) and *p*_*I*_(*t*) shared the same temporal profile except for a fixed relative delay. Both ramped linearly from 0 to 1 over a 20 ms period, maintained the steady-state value for the duration of the stimulus, and then decayed back to zero. The inhibitory delay, defined as the onset difference between *p*_*E*_(*t*) and *p*_*I*_(*t*), was varied between −2 and +2 ms. Firing rates were set to *r*_*E*_ = *r*_*I*_ = 50 Hz, and the numbers of afferents were *n*_*E*_ = *n*_*I*_ = 250. Peristimulus time histograms (PSTHs) of the reference neuron were computed from 100–1000 independent trials.

#### Transient stimuli

In this mode, each afferent fired a single action potential in response to a brief stimulus. Temporal jitter was introduced so that the afferent spikes did not arrive synchronously across inputs. The distribution of excitatory postsynaptic current (EPSC) arrival times to the reference and inhibitory neurons, denoted *h*_*E*_(*t*), was modeled as a Normal distribution scaled by *p*_*E*_*n*_*E*_:

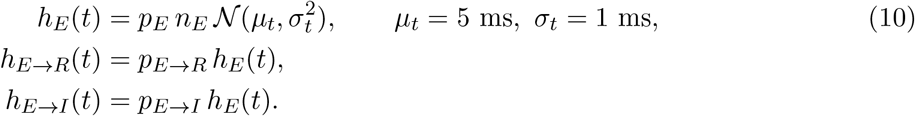

Here, *h*_*E*→*R*_(*t*) and *h*_*E*→*I*_(*t*) represent the effective distributions of EPSC arrival times in the reference and inhibitory neurons, respectively, accounting for the fact that not all afferent spikes evoke synaptic currents unless *p*_*E*→*R*_ or *p*_*E*→*I*_ equal 1.

Each histogram was convolved with the unitary synaptic kernel *a*_*E*_*u*_PSC_(*t*), which defines the time course of a single postsynaptic current, to obtain the compound excitatory current in the reference and inhibitory neurons. Independent realizations of *h*_*E*_(*t*) were generated for each inhibitory cell and their evoked spike times documented.

A corresponding histogram of inhibitory spike times, *h*_*I*_(*t*), was compiled from the responses of the inhibitory neurons. This histogram was scaled by *p*_*I*→*R*_ and convolved with *a*_*I*_ *u*_PSC_(*t*), which defines the time course of a unitary inhibitory synaptic current. The resulting current was then conditioned on excitation by multiplying by *p*_*E*_, and delivered to the reference neuron together with the excitatory current. Each simulation was repeated 1000–5000 times to compile the peristimulus spike histogram of the reference neuron (Fig. 6A, middle panel, gray).

Methods for calculating the time-dependent probabilities and predicted spike times are provided in sect. 0.2 of the Supplementary Material.

## S1 Supporting information

### 0.1 Model and parameters

**Figure S1:**
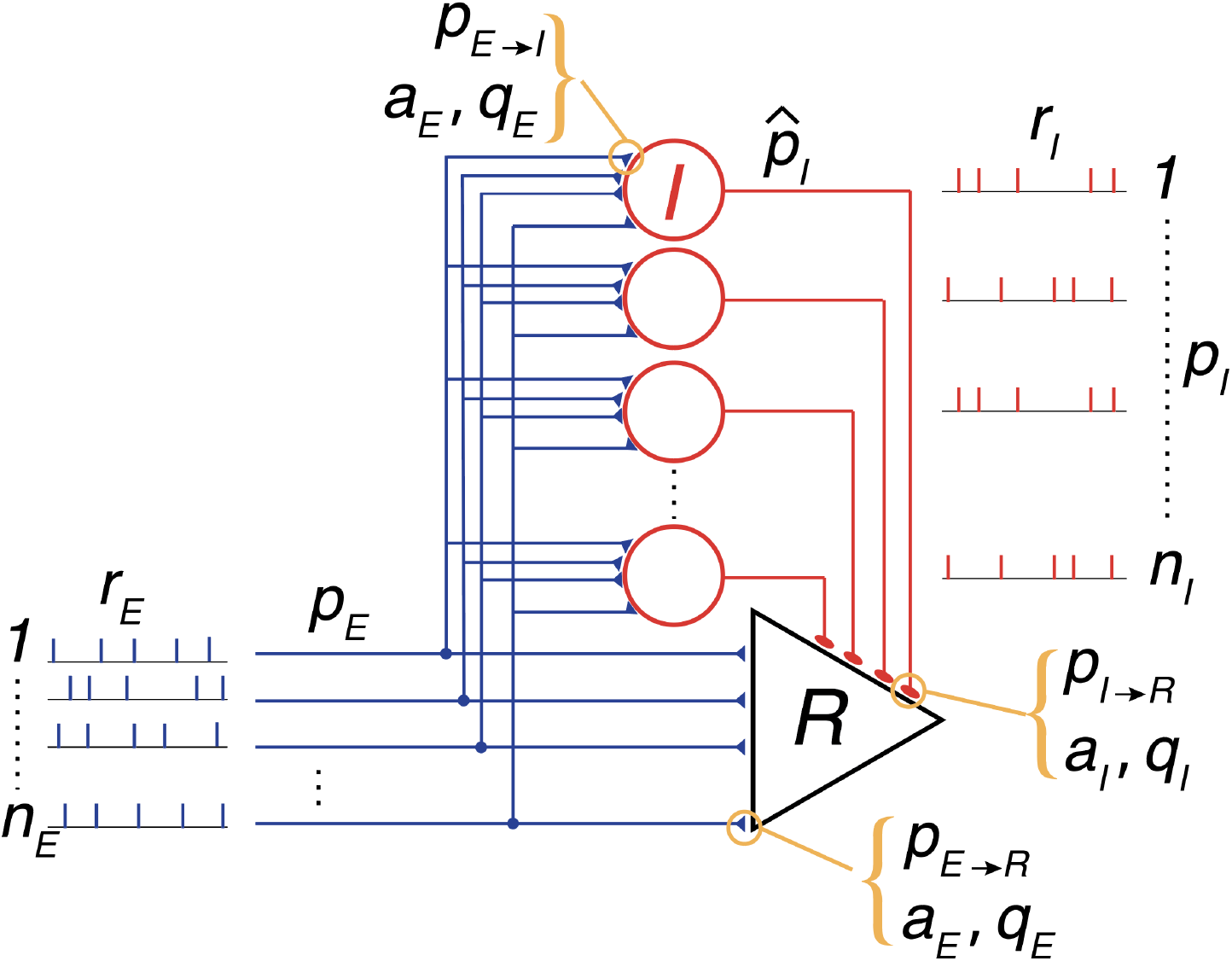
Model parameters.

The network is configured as feedforward inhibitory circuit, with a reference neuron (triangle, Fig. S1) and *n*_*I*_ inhibitory neurons, each receiving *n*_*E*_ excitatory inputs from an external source; the reference neuron also receives inputs from the local inhibitory neurons and is the main output of the network. In response to the stimulus, the external afferents fire either single (brief) action potential or trains (sustained) of action potentials. The model parameters are defined as follows.

**<u>Afferents:</u>**

*n*_*E*_ : number of afferents (100-250)

*p*_*E*_ : probability that an afferent will fire in response to a stimulus

*r*_*E*_ : the firing rate of afferents for long duration stimulus (set to 50 Hz)

*p*_*E*→*R*_ : the probability that an afferent spike will evoke an EPSP in the reference cell (synaptic efficacy). For simplicity, it is set to 1, indicating a reliable synapse.

*p*_*E*→*I*_: probability that an afferent spike will evoke an EPSP in an inhibitory cell (*p*_*E*→*I*_ = 1 if reliable synapse).

*a*_*E*_ : the amplitude of a single excitatory synaptic potential (250 µV) or current (10.3 picoAmps or pA) in both *I* cells and the reference cell.

*a*_*I*_ : the amplitude of a single inhibitory synaptic potential (-250 µV) or current (-10.3 pA) in the reference cell.

*q*_*E*_ : the charge transfer of a single excitatory synapse in both *I* cells and the reference cell (0.0557 picoCoulombs or pC).

*q*_*I*_ : the charge transfer (-0.0557 pC) of a single inhibitory synapse on the reference cell. Also expressed as fraction of excitatory charge transfer *q*_*I*_ = *k*_*q*_*q*_*E*_, *k*_*q*_ ∈ [0, 1]

*n*_*I*_ : number of afferents from inhibitory neurons (20-50). Also expressed as a fraction of the number of excitatory afferents: *n*_*I*_ = *k*_*n*_*n*_*E*_

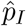: the probability that an *I* cell will fire during a stimulus

*q*_*I*_ : the charge transfer (-0.0557 pC) of a single inhibitory synapse on the reference cell. Also expressed as fraction of excitatory charge transfer *q*_*I*_ = *k*_*q*_*q*_*E*_, *k*_*q*_ ∈ [0, 1]

*r*_*I*_ : the firing rate of *I* neurons. Also expressed as *k*_*r*_*r*_*E*_, *k*_*r*_ ≥ 0

*p*_*I*→*R*_ :probability that a spike from an *I* neuron will evoke an IPSP in the reference cell (*p*_*I*→*R*_ = 1 if reliable synapse). Also expressed as a fraction of the excitatory synaptic efficacy: 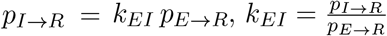

To simplify simulations,

1. *p*_*E*→*R*_ is set to 1 so that the probability of an EPSP appearing in the reference cell is just *p*_*E*_.
2. The effective probability (*p*_*I*_) of an IPSP appearing in the reference cell , accounting for the number, strength, and firing rate of inhibitory input relative to excitatory input (see below), is:

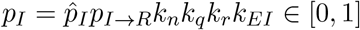

where 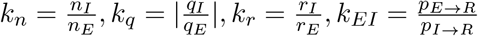 .

### 0.2 Calculating firing probability with sustained input

Many neurons, in response to sustained synaptic input, enter the oscillatory firing regime. Typically, the firing rate depends on the mean synaptic current (so-called firing rate-current, or f-I curve). During a sustained stimulus, the afferents and *I* neurons trains of EPSPs and IPSPs, respectively, producing a synaptic barrage in the postsynaptic neuron.

The first step is to express the net synaptic current in terms of *p*_*E*_ and *p*_*I*_. During a stimulus, each afferent is modeled as generating a Poisson spike train,

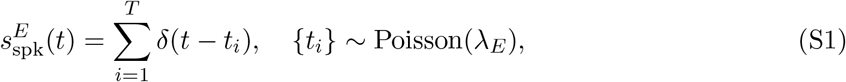

where *λ*_*E*_ denotes the rate of the homogeneous Poisson process. The excitatory postsynaptic current (EPSC) train is given by

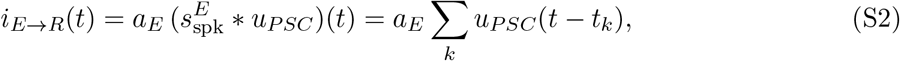

where ∗ denotes convolution. The kernel *u*_*PSC*_(*t*) represents the unitary EPSC generated by a single afferent and is normalized to unit peak, so that the amplitude *a*_*E*_ specifies the EPSC peak magnitude.

If there are *n*_*E*_ afferents, the total current is

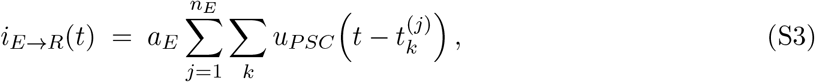

where 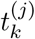 denotes the *k*-th spike time of afferent *j*.

If a stimulus causes each of the *n*_*E*_ excitatory afferents to fire with probability *p*_*E*_, and if each spike successfully generates an EPSC in the reference cell with probability *p*_*E*→*R*_, then the effective rate of excitatory events is

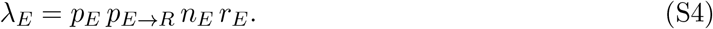

The corresponding mean excitatory current is therefore

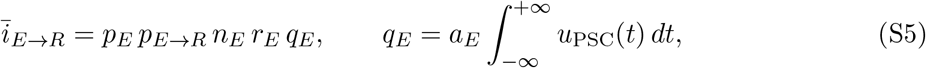

where *q*_*E*_ denotes the total charge transfer associated with a single EPSC, and *a*_*E*_ is a scaling factor converting the normalized postsynaptic current waveform *u*_PSC_(*t*) to physical units.

To calculate the inhibitory current into the reference cell during stimulation, the spike trains of the *I* neurons are first considered. The afferent inputs to the *I* neurons are analogous to those to the reference cell and yield the mean excitatory current

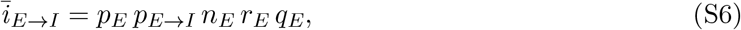

where *p*_*E*→*I*_ denotes the probability that an afferent spike evokes an EPSP in an *I* neuron. When this mean excitatory current exceeds the rheobase current *i*_*rh*_, each *I* neuron generates a spike train

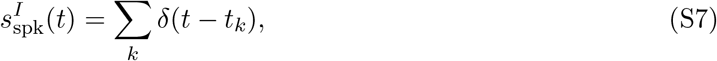

with firing rate *r*_*I*_, which depends on the input current and need not follow a Poisson process.

The inhibitory current generated in the reference cell by the population of inhibitory neurons is computed in several stages. First, the probability that an inhibitory neuron fires in response to a stimulus is defined as

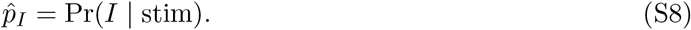

For simulations with an LIF model, 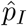was empirically measured as the probability that a stimulus evoked at least one spike during the stimulus interval (0.5–1 s). In principle, 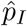 can also be expressed analytically as the probability that an inhibitory cell receives at least a threshold number of excitatory inputs within an integration time window.

If there are *n*_*I*_ inhibitory neurons firing asynchronously, and each spike generates an IPSC in the reference cell with probability *p*_*I*→*R*_, then the total inhibitory current is

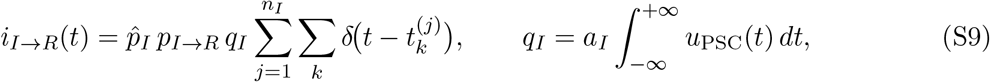

where *q*_*I*_ denotes the charge transfer associated with a single IPSC.

The mean inhibitory current is given by

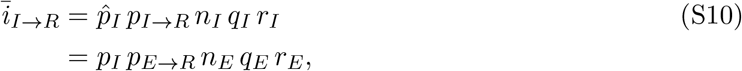

where *p*_*I*_ is the effective inhibitory probability that accounts for network size, synaptic efficacy, synaptic strength, and relative firing rate at the reference cell:

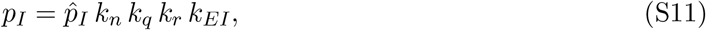

with 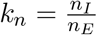 denoting the ratio of neuron numbers, 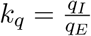 the ratio of charge transfers, 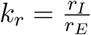 the ratio of firing rates, and 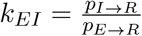 the ratio of synaptic efficacies.

Because of the feed-forward structure, the surviving excitatory current is

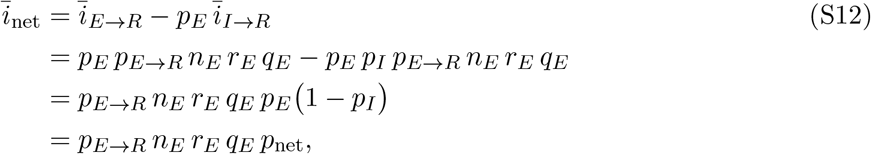

where the effective net excitation probability is defined as

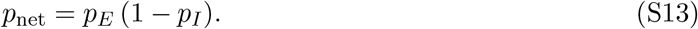

This expression represents the probability that an EPSP in the reference cell survives feed-forward inhibition recruited by the same presynaptic excitatory population.

In the sub-oscillatory (subthreshold) regime, firing may still occur due to voltage fluctuations that produce random threshold crossings. Let *X* denote the number of *surviving* (uncanceled) EPSPs received during the interval [*t*_0_, *t*_0_ + Δ*t*], so that *X* ∼ Binom(*n*_*E*_, *p*_net_), where *p*_net_ = *p*_*E*_ *p*_*E*→*R*_ (1 − *p*_*I*_). The fluctuation-driven firing probability is then

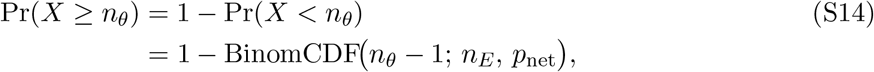

where *n*_*E*_ is the number of presynaptic excitatory afferents and *n*_*θ*_ is the minimum number of *uncanceled* EPSPs required to reach rheobase within Δ*t*.

The probability above is sigmoidal in *p*_net_ and approaches unity as the input approaches the oscillatory (tonic spiking) regime. To obtain a continuous transition between fluctuation-driven and oscillatory firing, define

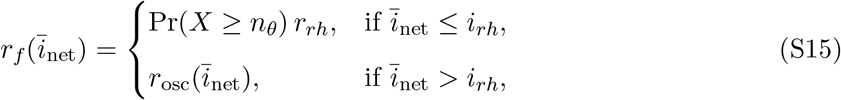

where

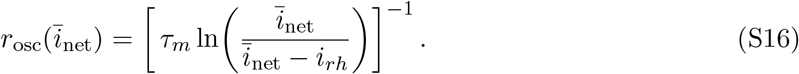

An alternative interpretation, under a Poisson point-process view, is that at rheobase the probability of at least one spike in a window of duration Δ*t* is

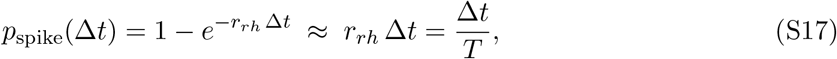

where *T* is the mean interspike interval at rheobase (*r*_*rh*_ = 1*/T*) and the approximation holds for Δ*t* ≪ *T* . Fluctuation-driven firing then scales this per-bin probability by the survival tail:

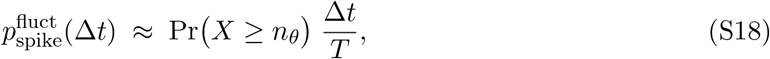

with *X* ∼ Binom(*n*_*E*_, *p*_net_) as defined above. Equivalently, dividing by Δ*t* yields the subthreshold rate

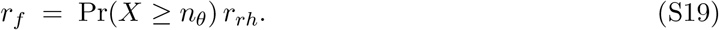

#### Calculating firing probability with transient input

In contrast to sustained stimulation, brief inputs produce a near-synchronous arrival of EPSPs and IPSPs, placing the neuron in a regime where spiking is highly sensitive to both the amplitude and precise timing of synaptic events. Consequently, the derivation and resulting expressions differ from those obtained under sustained input.

The analysis considers a reference neuron that receives *n*_*E*_ excitatory inputs from an external source and *n*_*I*_ inhibitory inputs from local interneurons. During stimulation, afferent EPSPs appear in both the reference cell and the local interneurons with probability *p*_*E*_. The probabilities that an afferent spike evokes an EPSP in the reference and *I* cells are denoted by *p*_*E*→*R*_ and *p*_*E*→*I*_, respectively. Thus, the probabilities of observing an EPSP in each target cell are

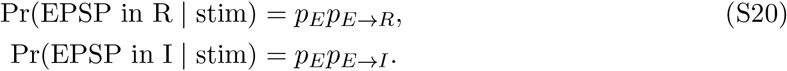

If each afferent fires a single action potential whose timing is Gaussian distributed during the stimulus, the expected compound EPSP evoked in the reference cell is given by:

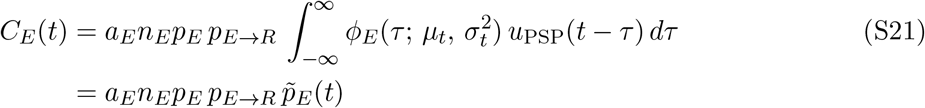

where 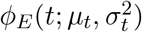 is the Gaussian probability density of presynaptic spike times, centered at *µ*_*t*_ with temporal dispersion *σ*_*t*_. The kernel *u*_PSP_(*t*) describes the time course of a unitary postsynaptic potential (e.g., an alpha function) with amplitude 1, *a*_*E*_ is the amplitude (in mV) of a single EPSP, and *n*_*E*_ is the number of afferents. Thus, 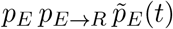 represents the instantaneous probability density that an excitatory input occurs at time *t*.

The arrival-time density *𝒫*_*I*_(*t*) of inhibition can be empirically estimated by constructing a histogram *h*_*I*_(*t*) from the activities of the *n*_*I*_ inhibitory neurons. It is expressed as

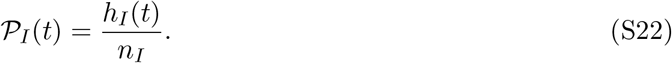

To isolate the timing of inhibition that is actually driven by the stimulus, the conditional inhibitory arrival-time density is computed only from stimulus presentations that successfully elicit excitatory input (and thus potential spiking) in the inhibitory neurons. A histogram *h*_*I*_(*t* | stim) of inhibitory spike times is constructed from these responsive trials and normalized by the number of inhibitory neurons *n*_*I*_ and the number of such stimulus repetitions *n*_stim_:

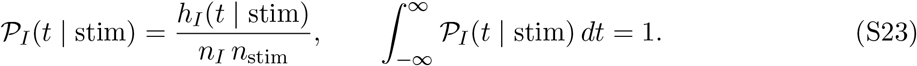

This conditional density therefore represents the temporal distribution of inhibitory activity given that the stimulus successfully activated the inhibitory population.

Then, the time-varying probability density of inhibition in the reference cell during stimulation is given by

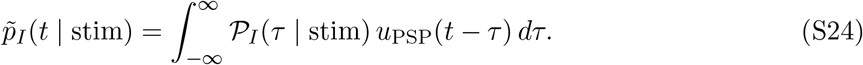

The compound IPSP can then be expressed, relative to the compound EPSP, as:

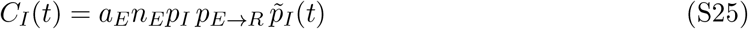

where *p*_*I*_ is an effective probability that incorporates factors related to the number of inhibitory neurons, synaptic strength, and synaptic efficacy:

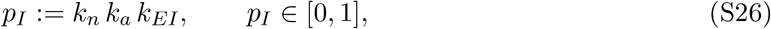

with 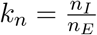 denoting the ratio of inhibitory to excitatory synaptic inputs, 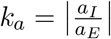 the ratio of inhibitory to excitatory amplitudes, and 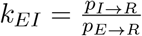 is the ratio of synaptic efficacies.

The corresponding conditional probability density of inhibition in the reference cell is then

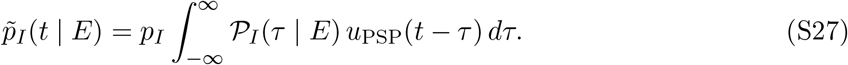

The net compound EPSP that is not canceled by IPSPs is given by

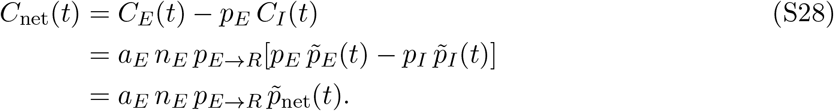

The net compound EPSP that is not canceled by IPSPs is given by

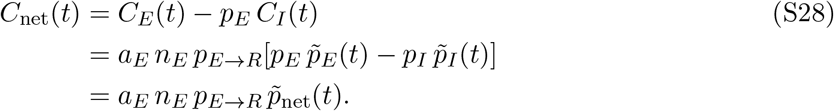

The time-varying factor 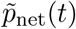 represents the instantaneous probability density of net excitation—that is, the component of the excitatory drive that is not canceled by inhibition. Because the difference 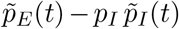can become negative (for example, when the compound IPSPs peak after a delay corresponding to the falling phase of the compound EPSP), a lower bound of 0 is imposed so that this term remains interpretable as a probability density.

Moreover, because inhibition in a feedforward circuit co-varies with excitation and both vanish in the absence of a stimulus, 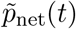 can be equivalently expressed as

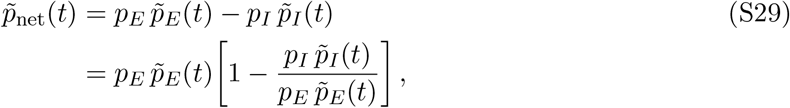

where the ratio 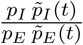 quantifies the instantaneous relative strength of inhibition to excitation.

The probability that the cell fires at time *t* is the probability that the membrane potential exceeds the threshold *v*_*θ*_. Equivalently, this is the probability that the net number of excitatory postsynaptic potentials (EPSPs) surviving inhibition exceeds the minimum number *n*_*θ*_ required to reach threshold. Let *X*(*t*) denote the number of uncanceled EPSPs received during the interval [*t, t* + Δ*t*]. The firing probability is then given by the upper tail of a binomial distribution:

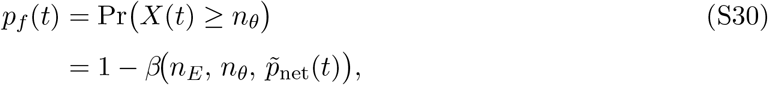

where 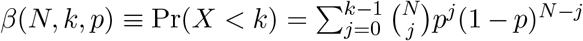 is the cumulative distribution function of the binomial distribution with *N* trials and success probability *p*. The green curve in Fig. 6A (bottom panel, main text) illustrates this firing probability.

Finally, the probability that an action potential occurs at a specific time *t*_*i*_ is given by the probability of firing at *t*_*i*_, multiplied by the probability of not having fired at any earlier time:

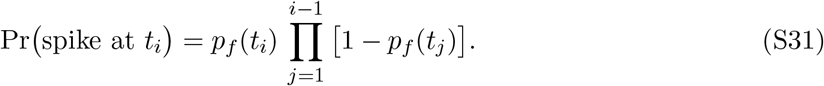

This expression corresponds to the magenta curve in Fig. 6A (middle panel).

#### Correcting for conductance

**Figure S2:**
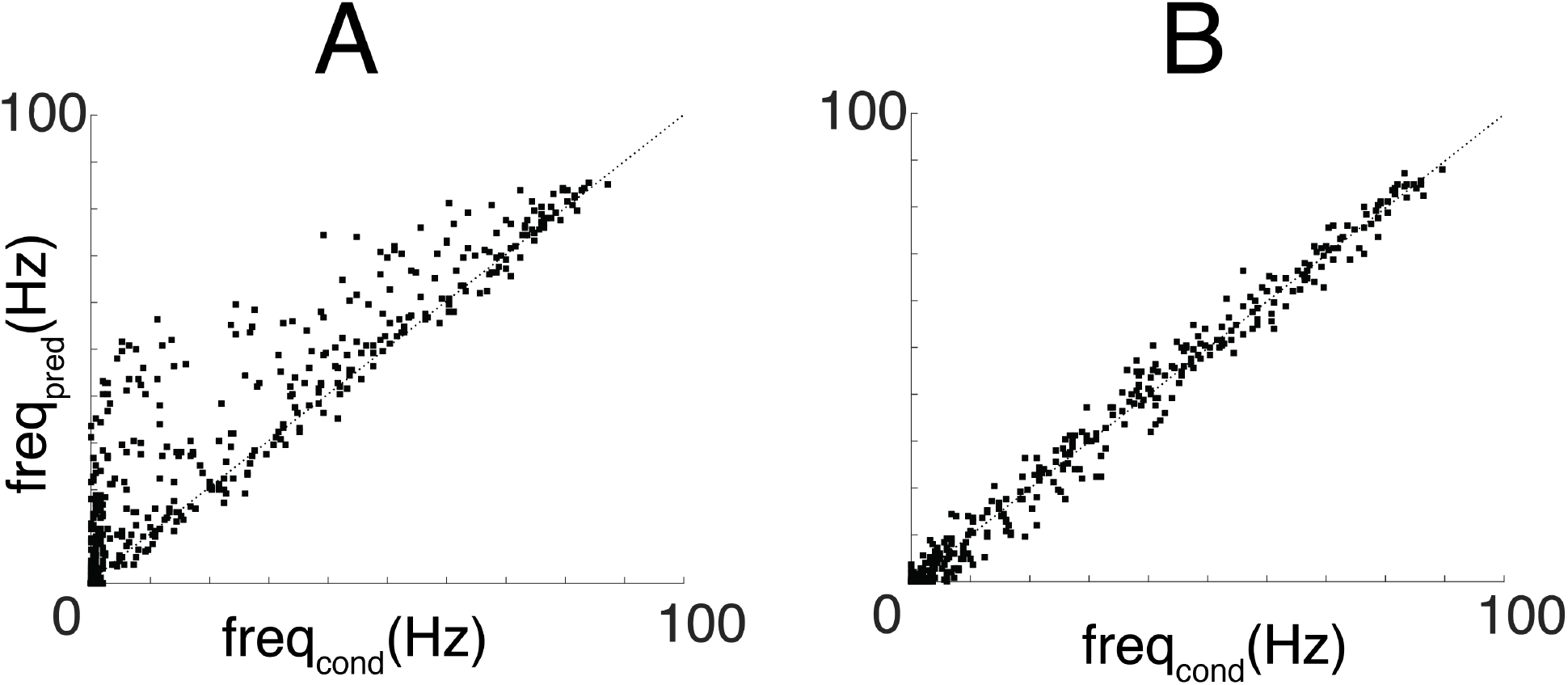
Effects of conductance. **A**, plot of firing rate predicted with with EPSPs with probability *p*_*net*_ vs firing rate evoked with conductance-based excitatory and inhibitory EPSPs and IPSPs. **B**, same but with predicted frequency corrected for voltage-dependent non-linear summation of PSPs.

The voltage-dependent unitary postsynaptic currents for excitation and inhibition (in nA) were defined as

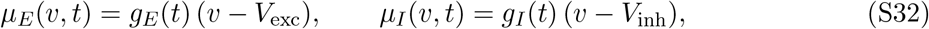

with corresponding charge transfers

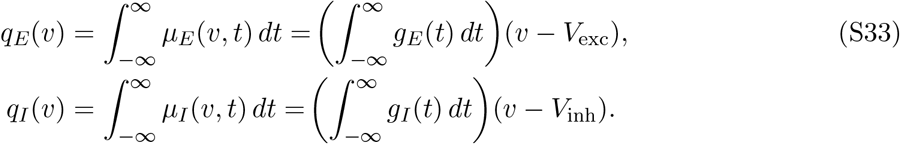

Here, the charge transfer *q*(*v*) has units of picoCoulombs (pC), and the conductances *g*_*E*_(*t*) and *g*_*I*_(*t*) have units of nanoSiemens (nS). Each conductance followed an alpha function:

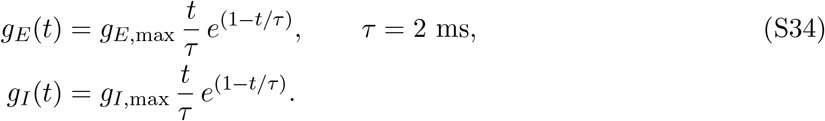

The mean voltage-dependent excitatory and inhibitory currents are

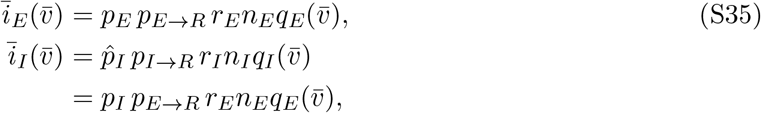

where 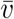 denotes the mean membrane potential under synaptic input in the absence of spiking, 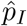is the probability that a stimulus evokes firing in an *I* cell (see above), and *p*_*I*_ is the effective inhibitory probability given by

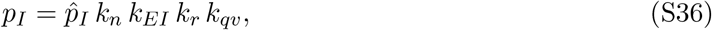

with

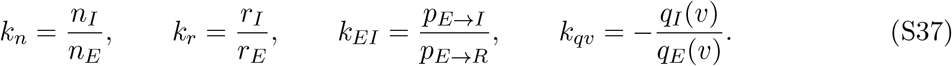

Thus, the net current is given by

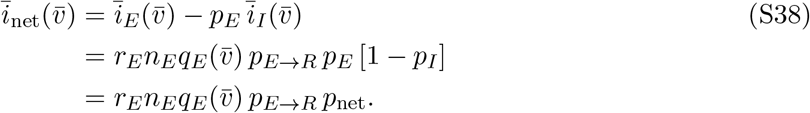

To examine how nonlinear summation affects predicted firing, the simulations described in the main text (Methods) were repeated, but modeled synaptic inputs as conductances rather than currents. To produce equal amplitudes (+250 *µ*V for EPSPs and −250 *µ*V for IPSPs) at the resting potential (−70 mV), with excitatory and inhibitory reversal potentials of 0 and −80 mV, respectively, the maximum inhibitory conductance (*g*_*Imax*_ = 1.045 nS) was set to be seven times larger than the excitatory conductance (*g*_*Emax*_ = 0.147 nS). The firing rate in this conductance-based mode of the simulation is denoted *f*_cond_.

The predicted firing rate *f*_pred_ was obtained as follows. First, the mean evoked membrane potential 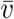 was estimated by repeating the simulations above with action potentials disabled. Second, a Poisson spike train was generated with rate *λ* = *n*_*E*_*p*_net_*r*_*E*_ (with *p*_*E*→*R*_ = 1). Third, the net synaptic current was obtained by convolving this spike train with a peak-normalized alpha function scaled by the excitatory conductance amplitude (*a*_*E*_; see Methods). Finally, the resulting current was delivered to the LIF neuron, and the firing rate was measured.

The firing of the reference cell in response to input depends on the parameters listed above (Model and Parameters) and in Table 2 of the Methods, summarized compactly in Eq. S38.

To evaluate the effectiveness of the conductance correction term (Eq. S38), simulations were performed using random values for selected parameters (*p*_*E*_, *p*_*E*→*I*_, and *k*_*n*_), while keeping the others fixed. As noted in the main text, changes in *p*_*E*→*I*_ affected the firing rate *r*_*I*_ (and hence *k*_*r*_) of inhibitory neurons and altered the I–O curves. Similarly, variations in *k*_*n*_ influenced the magnitude of inhibition to the reference cell and its I–O relation. Accordingly, 1000 trials were run with random combinations of *p*_*E*_ (0.6–1; 0.6 being the minimum required to evoke firing), *p*_*E*→*I*_ (0.5–1), and the inhibitory–excitatory ratio *k*_*n*_ (0.2–1), generating a wide range of firing frequencies. Plotting *f*_cond_ against *f*_pred_ (Fig. S2A) showed that *f*_pred_ systematically overestimated *f*_cond_, as many points lay above the unity-slope line (dotted). Incorporating the conductance correction substantially improved the correspondence (Fig. S2B).

#### Generation of bimodal tuning curves

**Figure S3:**
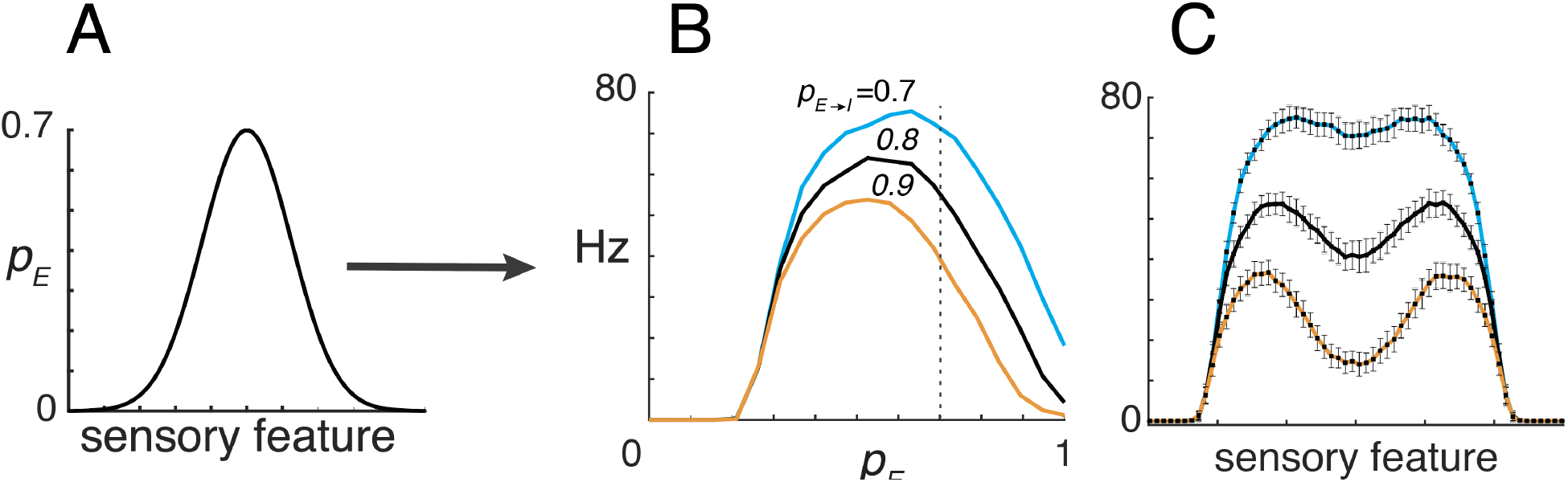
Effects of large tuned input. **A**, tuned input with peak of 0.7. **B**, I-O curves obtained when *π*_*E*→*I*_ increased with *p*_*E*_ set at at 0.7, 0.8, and 0.9. **C**, tuned responses showed a dip at the center .

In Fig. 4D of the main text, simulations were performed with small amplitude tuning curve (peak =0.35) under the condition that *π*_*E*→*I*_(= *p*_*E*_*p*_*E*→*I*_) increased linearly with *p*_*E*_ with fixed values of *p*_*E*→*I*_ . With this amplitude, the firing fell on the rising edge of the I-O curves and multiplicative gain modulation was produced. In Fig.S3, the amplitude of the tuned input was doubled to 0.7 (**A**). Because the evoked firing corresponded to the decaying edge of the I-O curves (**B**), the tuned responses (**C**) had a dip at the center.

#### Effects of conductance for brief stimuli

**Figure S4:**
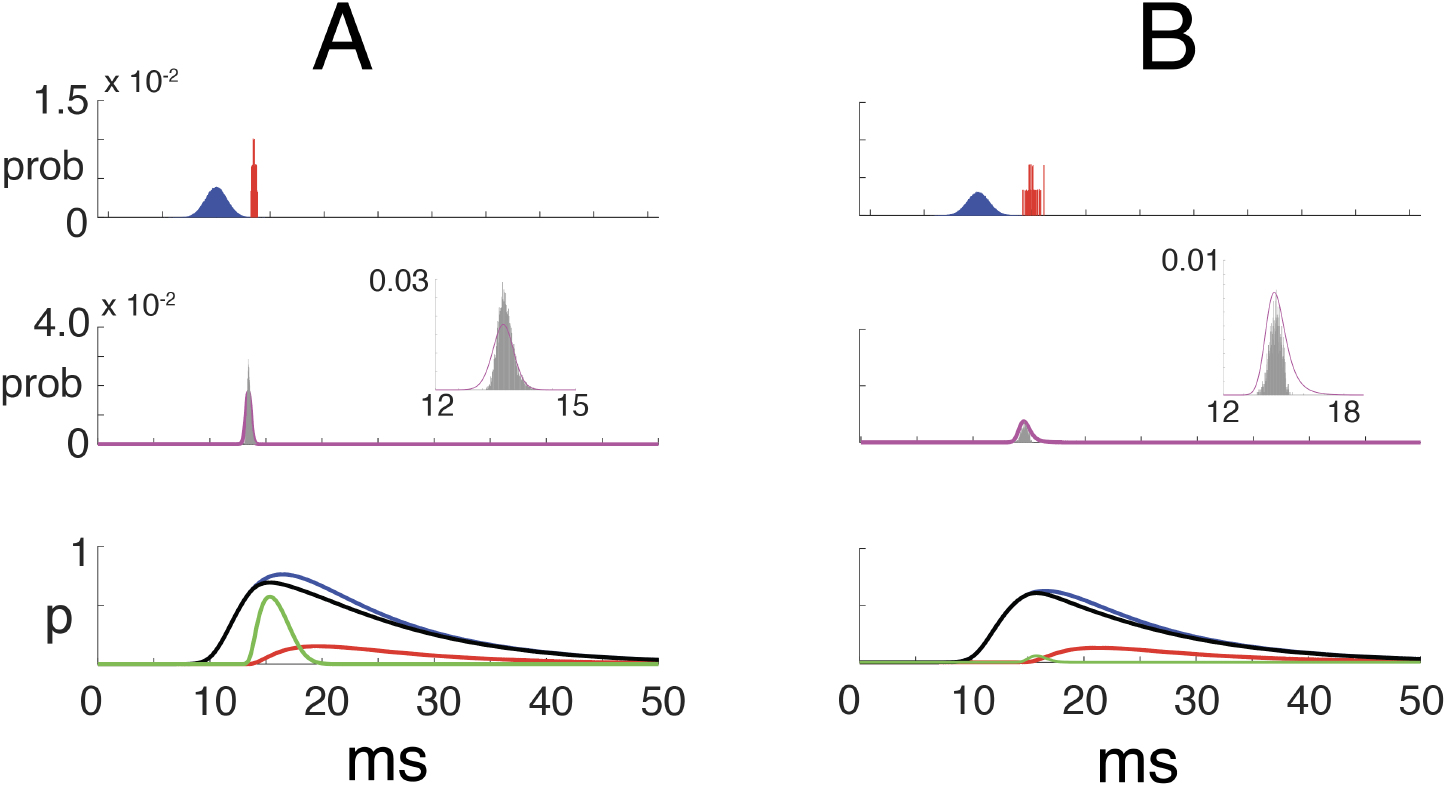
Predicting firing probability for transient stimuli in conductance mode. **A**,**top**, probability distributions of arrival times of EPSP (blue) and IPSP (red) in the reference cell. **Middle**, predicted firing probability (magenta) overlaid on the spike time histogram (gray); **Inset**, magnified view. **Bottom**, superimposed traces of 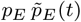 (blue), 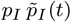 (red), net input 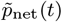(black), and threshold-crossing probability (green). *p*_*E*_ = 0.95, *k*_*n*_ = 0.2, *k*_*a*_ = 1, *k*_*EI*_ = 1. **B**, same but with *p*_*E*_ = 0.75 . **Model parameters:** *n*_*E*_ = 100, *n*_*I*_ = *k*_*n*_ *n*_*E*_ = 20, *g*_*E*_ = 0.147 nS, *g*_*I*_ = 1.045 nS, Histograms compiled from 5,000 trials with Bin width = 0.01 ms

#### Simulations with brief stimuli and synaptic conductances

The simulations conducted in the main text with transient stimuli assumed linear summation of synaptic inputs. However, there are scenarios where summation may be sublinear, such as when synapses are in close proximity or when inhibition causes significant shunting. To assess whether the model holds under these conditions, simulations were performed using conductance-based synaptic inputs (see Methods). Figure S3 shows that the results were qualitatively similar to those from the linear model (compare with Fig. 6 in the main text). The effects of conductance were not more pronounced, partly because action potentials occur primarily on the rising edge of 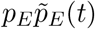 (compare the middle and bottom traces), before significant overlap with *p*_*I*_(*t*) occurs.

